# Aberrant phase separation is a common killing strategy of positively charged peptides in biology and human disease

**DOI:** 10.1101/2023.03.09.531820

**Authors:** Steven Boeynaems, X. Rosa Ma, Vivian Yeong, Garrett M. Ginell, Jian-Hua Chen, Jacob A. Blum, Lisa Nakayama, Anushka Sanyal, Adam Briner, Delphi Van Haver, Jarne Pauwels, Axel Ekman, H. Broder Schmidt, Kousik Sundararajan, Lucas Porta, Keren Lasker, Carolyn Larabell, Mirian A. F. Hayashi, Anshul Kundaje, Francis Impens, Allie Obermeyer, Alex S. Holehouse, Aaron D. Gitler

**Affiliations:** Department of Molecular and Human Genetics, Baylor College of Medicine, Houston, TX 77030, USA; Jan and Dan Duncan Neurological Research Institute, Texas Children’s Hospital, Houston, TX 77030, USA; Therapeutic Innovation Center (THINC), Baylor College of Medicine, Houston, TX 77030, USA; Center for Alzheimer’s and Neurodegenerative Diseases (CAND), Texas Children’s Hospital, Houston, TX 77030, USA; Dan L Duncan Comprehensive Cancer Center (DLDCCC), Baylor College of Medicine, Houston, TX 77030, USA; Department of Genetics, Stanford University School of Medicine, Stanford, CA 94305, USA; Department of Chemical Engineering, Columbia University, New York, NY, 10027, USA; Department of Biochemistry and Molecular Biophysics, Washington University School of Medicine, St. Louis, MO 63110, USA; Center for Biomolecular Condensates, Washington University in St Louis, St. Louis, MO 63130, USA; Molecular Biophysics and Integrated Bioimaging Division, Lawrence Berkeley National Laboratory, Berkeley, CA 94720, USA; Department of Anatomy, University of California, San Francisco, CA 94143, USA; Clem Jones Centre for Ageing Dementia Research (CJCADR), Queensland Brain Institute (QBI), The University of Queensland, Brisbane, QLD 4072, Australia; VIB-UGent Center for Medical Biotechnology, 9000 Gent, Belgium; VIB Proteomics Core, 9000 Gent, Belgium; Department of Biochemistry, Ghent University, 9000 Gent, Belgium; Department of Biochemistry, Stanford University School of Medicine, Stanford, CA 94305, USA; Department of Pharmacology, Escola Paulista de Medicina (EPM), Universidade Federal de São Paulo (UNIFESP), Sao Paulo, Brazil.; Department of Integrative Structural and Computational Biology, The Scripps Research Institute, La Jolla, CA 92037, USA; Department of Computer Science, Stanford University, Stanford, CA 94305, USA; Chan Zuckerberg Biohub, San Francisco, CA 94158, USA

**Keywords:** Phase separation, cathelicidin, protein aggregation, membraneless organelles, frontotemporal dementia, amyotrophic lateral sclerosis, antimicrobial peptide, crotamine, RAN translation, LL-37

## Abstract

Positively charged repeat peptides are emerging as key players in neurodegenerative diseases. These peptides can perturb diverse cellular pathways but a unifying framework for how such promiscuous toxicity arises has remained elusive. We used mass-spectrometry-based proteomics to define the protein targets of these neurotoxic peptides and found that they all share similar sequence features that drive their aberrant condensation with these positively charged peptides. We trained a machine learning algorithm to detect such sequence features and unexpectedly discovered that this mode of toxicity is not limited to human repeat expansion disorders but has evolved countless times across the tree of life in the form of cationic antimicrobial and venom peptides. We demonstrate that an excess in positive charge is necessary and sufficient for this killer activity, which we name ‘polycation poisoning’. These findings reveal an ancient and conserved mechanism and inform ways to leverage its design rules for new generations of bioactive peptides.

## INTRODUCTION

Nucleotide repeat expansions are implicated in an expanding group of human diseases^1,2^. There are several different subclasses. Polyglutamine (polyQ) diseases, at least 9 different diseases each caused by the expansion of an exonic CAG repeat, are perhaps the most well-known^3^. These disease genes harboring the CAG repeat expansions get translated into proteins with an expanded polyQ tract, which makes them aggregation-prone, resulting in the formation of inclusion bodies as the hallmark pathology. Besides polyQ, expansions of exonic GCG and CGG repeats produce aggregation-prone polyalanine and polyglycine proteins, and cause a series of other diseases^4,5^. Repeat expansions in non-coding parts of genes can also cause disease. For example, in Fragile X-syndrome, large GC-rich expansions within promoter or 5’ untranslated regions (UTR) become hypermethylated, inhibiting transcription and resulting in haploinsufficiency^6^. When expanded nucleotide repeats are transcribed into RNA, the repeat RNAs themselves can be toxic–for example, CUG/CCUG intronic/3’UTR RNA repeats in myotonic dystrophies^7^. These repeat RNAs have a propensity to condense and can trap important RNA-binding proteins in the process, impairing their function. Even though, these distinctions between coding and non-coding repeats seemed simple at first, the picture has become more complicated. The repeat RNA of the exonic CAG repeats can be just as toxic as the intronic CUG ones^8^. Expanded repeats also can generate antisense transcripts, implicating additional types of repeat RNAs and proteins^9^. Moreover, repeat RNAs (even non-coding ones) can be aberrantly translated in an AUG-independent manner (referred to as RAN translation)^10^, greatly expanding the range of proteins that can be encoded from these nucleotide repeats.

An archetype of this complexity is the repeat expansion in the *C9orf72* gene. A hexanucleotide GGGGCC expansion in the first intron of a previously undescribed gene was found to be the leading genetic cause of amyotrophic lateral sclerosis (ALS) and frontotemporal dementia (FTD)^11,12^. Upon the discovery of this repeat expansion, an immediate question was one of mechanism—how does this expansion drive human disease? Evidence has emerged linking all of the aforementioned mechanisms to C9orf72 pathogenesis. Hypermethylation of the expanded repeat affects transcription of the gene^13^, which seems to be implicated in autophagy^14^, vesicle trafficking^15^ and immune function^16^. Repeat RNA foci (both sense and antisense versions) accumulate in patient cells^12,17^. RAN translation of the sense and antisense repeat generates so-called dipeptide repeats (DPRs) that are detected in the brain and spinal cord of patients harboring this mutation^17–19^. These are poly-GA, poly-GP, poly-PA, poly-GR and poly-PR (abbreviated as their amino acid codes from here on). GP and PA do not seem to be toxic, whereas GA has mixed effects based on the model being used. The two arginine-rich DPRs, GR and PR, are potently toxic in diverse model systems—ranging from yeast to flies and worms, to vertebrates and iPSC-derived motor neurons^20–23^.

Because GR and PR seem to be so toxic, there have been many different studies investigating the mechanisms of this toxicity. These studies have implicated disturbances in nucleocytoplasmic transport^24–26^, defects in stress granules^27,28^, mitochondrial defects^29^, perturbation of axonal transport^30^, proteasome deficits^31^, and inhibition of translation^28,32^. All of these mechanisms could help explain the etiology of the disease, which is undoubtedly complex, but we wondered how can two amino acids repeated head to tail cause such widespread cellular defects? In other words, why are PR and GR such promiscuous killers that seemingly hit many unrelated biochemical pathways at once?

Since their discovery a decade ago, the apparent “aspecificity” of PR and GR toxicity has confused the field. A unifying framework to explain the biological connection between their myriad effects and the observed target promiscuity has remained elusive. Here, we unexpectedly discover that the interactome of GR and PR is actually encoded in their biophysical behavior. We and others previously showed that PR and GR can undergo biomolecular condensation *in vitro* and target biomolecular condensates in cells^22,27,28,33^. Using new unbiased proteomics studies, we find that even the non-condensate effects of PR and GR can be explained by their “sticky” multivalency. Moreover, we find that nucleic acids are key modulators of their target space (i.e., the set of proteins that interact with GR and PR), explaining previously unresolved discrepancies between nuclear and cytoplasmic effects, and between disease models and patient material. Further, the recent identification of arginine-rich repeat peptides in at least three other degenerative conditions points at the emergence of a new class of repeat disorders centering around cationic peptides^34–36^, with potentially a shared pathophysiology.

Since our framework was able to connect pathological condensation events in a divergent set of biological pathways, we wondered whether we could harness it to predict novel physiological condensate proteins. Using machine learning approaches, we discovered that the promiscuous condensation-based toxicity of PR and GR is not limited to human disease but is an integral part of ancient natural anti-microbial defense systems and spider, scorpion, and rattlesnake venoms. Similar cationic peptides have evolved countless times independently throughout the tree of life, and all can drive aberrant condensation via electrostatic interactions. Finally, leveraging protein design, we show that a surplus of cationic charge is sufficient to turn an inert protein into a toxic killer peptide. Thus, a mechanism we name *polycation poisoning* is emerging as a powerful and unifying framework that guides our understanding of promiscuous killer peptides in health and disease, and may aid in the design of new generations of therapeutic bioactive peptides.

## RESULTS

### RNA concentration tunes the target space of arginine-rich dipeptides

In addition to C9orf72 ALS/FTD, other diseases also exhibit the same or similar arginine-rich peptides (**Fig. 1A**). RAN translation of GGCCTG (NOP56) and AGAGGG (TAF1) repeats, linked to spinocerebellar ataxia type 36 (SCA36) and X-linked dystonia and parkinsonism (XDP), generate PR and GR DPRs, respectively^34,35^. QAGR tetrapeptide repeats are produced from the CCUG repeat in myotonic dystrophy type 2 (DM2)^36^. As we have shown previously for PR and GR, these QAGR peptides similarly condense *in vitro* when added to cell lysate (U2OS, **Fig. 1B**). We previously showed that PR and GR form liquid droplets in the test tube when combined with nucleic acids^27^, and these assemblies—or ones using other cationic peptides—have been since used as very simple *in vitro* models to study the biophysics of more complex RNA-centric liquid condensates in the cell^37–40^. Surprisingly, when we add these peptides to cell lysate, we observe the formation of irregular networked condensates (**Fig. 1C**, **Fig. S1A**)—more reminiscent of the pathological gel-like condensates found in neurodegenerative disease^41,42^. Adding additional RNA into the mixture (i.e., total yeast RNA), caused these assemblies to change morphology and acquire spherical shapes more similar to the liquid-like behavior of some physiological condensates (e.g., nuclear bodies like nucleoli^43^). Because the addition of RNA dramatically affected the appearance of these test tube assemblies, we wondered what this would mean for their protein composition. As expected, its negative charge allows RNA to compete with proteins in the cell lysate for binding to the positively charged PR peptide (**Fig. 1D**, **Fig. S1B-D**). Using unlabeled quantitative mass spectrometry, we dissected what this competition looks like at the level of individual proteins (**Table S1**). We calculated the relative solubility of proteins in the test tube after spinning down the PR-induced condensates—as a measure of condensate partitioning— and clustered proteins according to their respective solubility profile as a function of the added RNA concentration (**Fig. 1E-F**). Around 50% of proteins in the mixture never partitioned into PR condensates and remained soluble throughout the experiment (cluster 1; *grey* proteins). For those proteins that condense with PR, 25% (clusters 2-5; *red* proteins) were outcompeted by low concentrations of RNA; the other 25% (clusters 6-9; *blue* proteins) were only outcompeted by high RNA concentrations or partitioned more strongly into the pellet upon the addition of RNA. Of note, this blue behavior is strikingly similar to the RNA-dependent re-entrant condensation behavior that has been observed for several RNA-binding proteins^38,44,45^. Thus, these results demonstrate that PR-interacting proteins can have distinct RNA-dependent interaction profiles. Interestingly, these three clusters—simply based on the *in vitro* condensation behavior of proteins with PR and RNA—each specifically enriched different functional categories of proteins (**Fig. 1G**). The blue class of proteins included canonical members of physiological condensates (e.g., nuclear speckles, nucleoli) or cellular assemblies that have been proposed to form via similar biophysical principles (i.e., heterochromatin^46,47^) (**Fig. S2**). Red proteins consisted of proteins not typically associated with physiological condensates but that have been implicated in protein aggregation or its modulation (**Fig. 1H**, **Fig. S1E-H**). Consistent with these findings, several of these distinct biological pathways—both red and blue—have been directly implicated in the toxicity observed in C9orf72 disease models and/or patients. These include nuclear import receptors^24,48^, protein arginine methyl transferases^24,49,50^, microtubules^30^, the proteasome^31^, the translational machinery^28,32, 51–53^, splicing factors^28,54^, nucleoli^27,28,55,56^, and heterochromatin^57^, among others. Thus, our simple test tube model recapitulates the target promiscuity of arginine-rich DPRs and provides a framework to understand how local RNA concentration modulates this target space, with important implications for pathophysiology (see **Discussion**).

**Figure 1:**
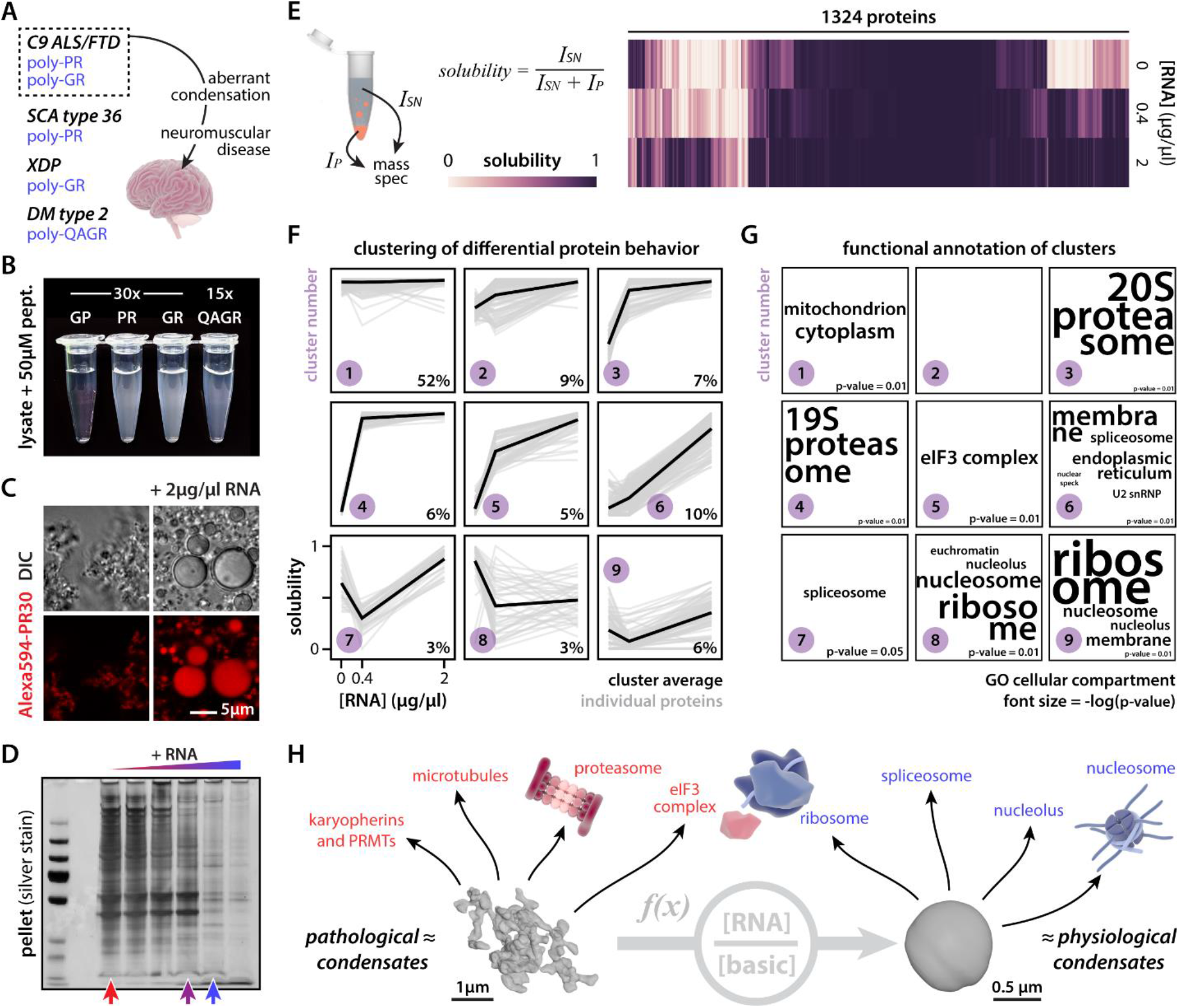
RNA modulates protein composition of *ex vivo* PR30 condensates. (A) Scheme indicating toxicity of cationic RAN peptides. (B) Cationic RAN peptides drive condensation of U2OS cell lysate. Heatmap showing quantification of RNA-dependent solubility for 1324 proteins. (C) Morphology or PR30 condensates is modulated by exogenous RNA addition (2 µg/µl RNA). (D) Silver stain indicating differential protein composition of PR30-induced condensates in response to RNA addition. (E) Heatmap showing quantification of RNA-dependent solubility for 1324 proteins. (F) Proteins cluster according to solubility responses as a function of RNA concentration. % indicates fraction of total proteins in the cluster. (G) Clusters differentially enrich for GO terms relating to subcellular compartments. Font size = -log(p-value). (H) Overview indicating that the morphology and composition of PR30 condensates are a function of the ratio of RNA concentration over basic protein concentration. All of the highlighted protein assemblies and factors have been directly implicated in C9orf72 ALS/FTD pathogenesis. 3D reconstructions of PR condensates were obtained via soft x-ray tomography. See also **Fig. S1**.

### RNA-dependent behavior is encoded in sequence and follows an electrostatics framework

PR and GR’s apparent target promiscuity has remained enigmatic; how are they able to target such a wide range of unrelated biological pathways? Because we now have in hand an *in vitro* model of this target space, we could ask whether this behavior and its modulation by RNA are encoded by a protein’s sequence. First, overlapping our data with a set of experimentally validated RNA-binding proteins^58^ showed that, as expected, blue proteins are enriched for RNA-binding activity compared to red and grey proteins (**Fig. 2A**). Second, based on amino acid composition, blue and red proteins separate based on their charge (**Fig. 2A-B**); blue proteins are on average more basic, red ones more acidic, and grey ones more neutral. We can even observe this effect when looking within protein families. 60S ribosomal proteins are on average more positively charged than the ones making up the 40S subunit, and this correlated with lower solubility profiles in our mass spectrometry experiment (**Fig. 2C**).

**Figure 2:**
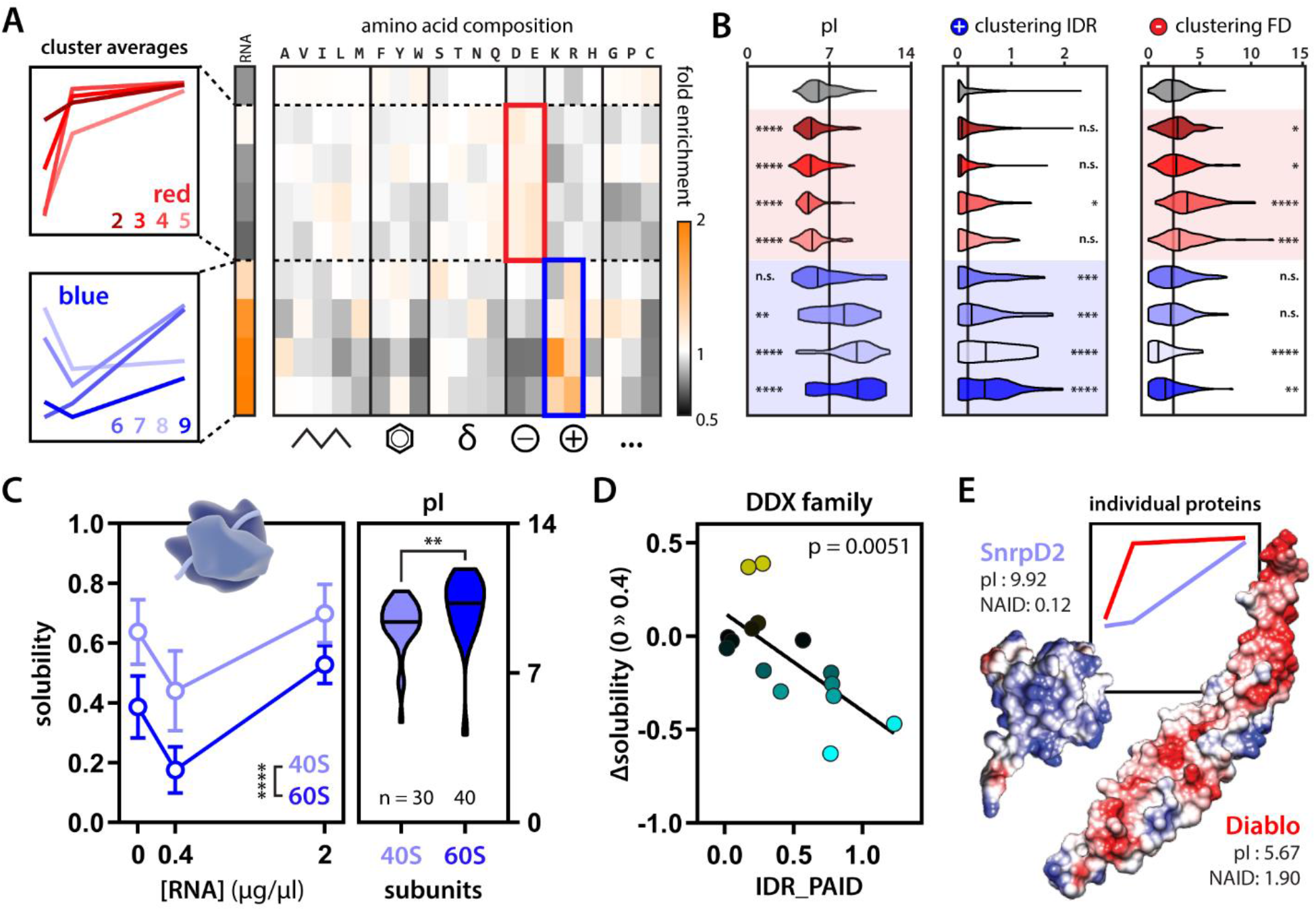
Electrostatic charge is a sequence feature modulating PR30 condensate partitioning. (A) Blue proteins are enriched for experimentally validated RNA-binding proteins and cationic amino acids. Red proteins have higher acidic amino acid composition. Grey proteins are enriched for neither. (B) On average, red proteins are more acidic and blue proteins are more basic compared to grey proteins (left). Moreover, on average, blue proteins have more clusters of basic residues in their IDRs (middle), while red proteins have more clusters of acidic residues in their folded domains (right) compared to grey proteins Kruskal-Wallis. (C) 40S and 60S ribosomal subunits show differential solubility profiles that correlate with changes in average pI. Two-way ANOVA and Mann-Whitney. (D) Positive charge clustering on the IDRs of DDX helicases correlates with their partitioning response to RNA (change in solubility upon addition of 0.4 µg/µl RNA. Linear regression. (E) Two example folded proteins with mirrored convex-concave solubility curves show opposite surface charge clustering (NAID = negative average inverse distance) and pI. Diablo (PDB: 1FEW)^108^, SnrpD2 (PDB: 1B34)^109^. See also **Fig. S3-4**. * p-value < 0.05, ** p-value < 0.01, *** p-value < 0.001, **** p-value < 0.0001.

While average charge can be informative, how charged residues are distributed across intrinsically disordered regions (IDRs) or on the surface of folded domains can substantially impact molecular interactions^59–62^. Therefore, we co-opted and extended our previously applied sequence clustering analysis^63^ to evaluate the position of acidic and basic residues in both IDRs and on the surface of neighboring fold domains (in linear space and three-dimensional space, respectively) (**Fig. S3**). By integrating proteome-wide structural predictions, this analysis revealed that—on average—blue proteins possess clusters of basic residues within IDRs, while red proteins possess clusters of acidic residues on the surface of folded domains (**Fig. 2B**, **Fig. S4**). We highlight two examples to further illustrate these observations. First, DEAD-box helicases (DDX) are a family of RNA chaperones that share the same type of folded domain typically flanked by IDRs. We identified red, blue, and grey DDX proteins in our assay, and, importantly, the positive charge clustering of their respective IDRs dictates their RNA-dependent condensation (**Fig. 2D, Fig. S1G-H**). Second, we identified two proteins that had virtually mirrored solubility profiles. SnrpD2 (a spliceosome component) shows very little acidic clustering on its folded domain but has a strongly basic pI, while Diablo (a mitochondrial protein) shows specific subregions with clusters of acidic residues (**Fig. 2E**). Despite having the same initial and final solubility in our mass spectrometry experiment, these proteins show opposite RNA-dependence, in line with their respective charge profiles. These analyses indicate that the differential protein-partitioning we observed in our mass spec experiment—on average—follows simple rules relating to the density and positioning of acidic and basic residues that tune protein-protein and protein-RNA interactions. We stress that these analyses report on average properties. Charge or the ability to bind RNA is predictive, but there are many examples of proteins that enrich differently. This argues that other—yet unresolved and probably more complex— sequence features and protein-protein interactions modulate protein partitioning in our assay.

### Generating a machine learning algorithm to predict protein condensation

Because the condensate partitioning behavior in our mass spectrometry experiment correlated with certain sequence features, we reasoned that we could in principle use our experimental data set to learn to predict such behavior *de novo*. We trained a machine-learning algorithm^64^ (see **Material & Methods**) to predict blue proteins from their red and grey counterparts (**Fig. 3A**). Next, we asked the algorithm to rank intracellular proteins based on their “color” (**Fig. 3B, Table S2**). As expected, top predicted blue proteins enriched for gene ontology terms associated with nuclear bodies, RNA and chromatin metabolism (**Fig. 3C**). Interestingly, the algorithm also enriched intermediate filament proteins, which have been shown previously to engage with arginine-rich DPRs via their disordered head domains^33^. None of these condensation-related gene ontology terms came up for the red and grey class of proteins nominated by the algorithm.

**Figure 3:**
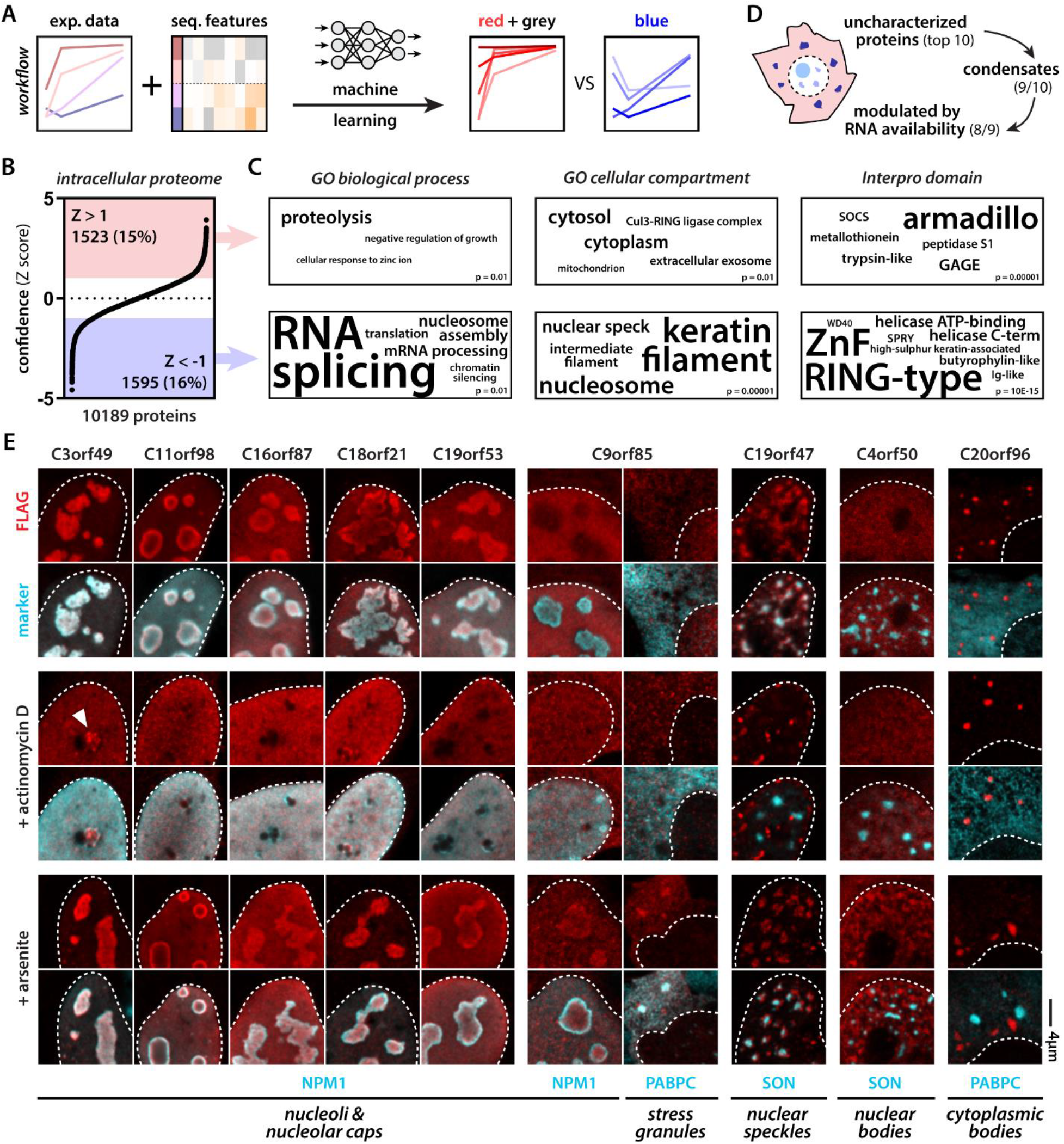
Machine learning predicts RNA-modulated phase separation of uncharacterized proteins. (A) Scheme illustrating workflow to predict red (concave) vs blue (convex) proteins. (B) The intracellular proteome ranked according to normalized confidence scores (Z score). (C) GO categories associated with predicted red and blue proteins. (D) Scheme highlighting breakdown of top 10 predicted blue unknown proteins. (E) 9 out of 10 predicted blue proteins target membraneless organelles. 8 do so in a manner that is dependent on nuclear or cytoplasmic RNA availability (2 µg/ml actinomycin D inhibits all transcription, 0.5 mM arsenite induces polysome disassembly and increases free cytoplasmic RNA levels). White arrowhead indicates nucleolar stress cap targeting of C3orf49. C9orf85 targets nucleoli and stress granules upon arsenite treatment. C19orf47 demixes from nuclear speckles upon transcription inhibition. C4orf50 condenses into nuclear bodies upon arsenite treatment. C20orf96 forms cytoplasmic bodies unresponsive to changes in RNA availability. C4orf54 binds the actin cytoskeleton (see **Fig. S5**).

Given blue proteins appear to consistently be associated with intracellular condensates, we decided to directly test our algorithm’s ability to predict condensation behavior for previously uncharacterized proteins. We tested the top ten uncharacterized proteins from our predicted blue protein list and expressed them in cells (**Fig. 3D, Table S2**). Seven of these formed or targeted condensates under normal conditions: C3orf49, C11orf98, C16orf87, C18orf21, and C19orf53 targeted nucleoli, C19orf47 localized to nuclear speckles, and C20orf96 formed unknown cytoplasmic condensates. This result clearly demonstrates that our machine-learning model can predict cellular and biophysical behavior. Since our initial mass spectrometry experiment investigated condensation as a function of RNA concentration, we wondered how changing RNA availability would influence the behavior of our seven condensate-forming proteins. To do this, we decided to stress cells in two ways to change nuclear and cytoplasmic RNA availability–transcriptional shutdown and polysome disassembly. First, we shut down all transcription with a high dose of actinomycin D^65^. This made four out of five of the nucleolar proteins demix from the nucleolus into the nucleoplasm—a well-known response of several nucleolar proteins^66,67^. C3orf49 also showed localization to the nucleolar periphery in assemblies reminiscent of nucleolar stress caps^68^. The nuclear speckle protein C19orf47 remained in a condensed state upon transcription inhibition, but demixed from the nuclear speckle, showing that its condensation behavior was also modulated by RNA. Second, stressing the cells with arsenite—known to cause polysome disassembly in the cytoplasm and rewire transcription in the nucleus^69^—made C9orf85 partition into stress granules. Peculiarly, in addition to stress granule partitioning, C9orf85 also translocated into nucleoli in parallel. C4orf50, previously found in the nucleoplasm, now formed abundant small nuclear granules that were clearly distinct from nuclear speckles. Transcription inhibition or arsenite stress did not affect C20orf96 condensates. Finally, C4orf54 localized to the actin cytoskeleton under all tested conditions (**Fig. S4**).

Combined, our algorithm predicted nine out of ten proteins correctly as condensate proteins, of which eight did so in a manner that was modulated by RNA availability. The only “incorrect” prediction, turns out to be a novel actin-binding protein. Interestingly, we and others have previously found that PR and GR can interact with actin^50^, microtubules^30^, and intermediate filaments^33^—highlighting that there may be more generally shared sequence features between cytoskeleton-binding and condensate proteins. Future studies will be required to investigate the functions of these new proteins and the functional significance of their condensate formation. In all, these findings indicate that our algorithm can predict condensate formation properties of proteins with high accuracy, allowing us and others to start systematically interrogating diverse proteomes.

### Condensation is a novel mode of action of bioactive killer peptides

In the above analysis, we restricted our search for novel condensate proteins to intracellular proteins since biomolecular condensates have been almost exclusively studied within the confines of the plasma membrane. But given the accuracy of our prediction tool, we reasoned that we could explore the possible existence of condensation events outside of the cellular context. When we fed sequences of secreted proteins into the algorithm, we unexpectedly found that it predicted several antimicrobial peptides (AMPs) to condense (**Fig. 4A, Table S2**). Animals, including humans, have evolved a wide array of AMPs^70^. A well-studied example is LL-37 (**Fig. 4B**). Some cationic AMPs have been shown to kill bacteria by permeabilizing their membranes, but for LL-37 and several other AMPs this does not seem to be their main mode of action^71^. Because of their positive charge, such peptides are expected to interact with membranes, hence, potentially crossing them. Indeed, cationic peptides are often called cell-penetrant since they are spontaneously taken up by both prokaryote and eukaryote cells^72^. This raised the question: what would happen once a cationic peptide like LL-37 enters the bacterial cytoplasm? To answer this, we first performed a similar *ex vivo* lysate assay as described above. Adding LL-37, or PR as a positive control, to *E. coli* lysate was able to potently drive condensate formation (**Fig. 4C**). Second, we treated live bacteria with LL-37 at a concentration corresponding to its minimum inhibitory concentration against a range of pathogens^73^. We subsequently imaged these bacteria using soft x-ray tomography, allowing us to gain information on their subcellular organization (**Fig. 4D**, **Fig. S5A**). LL-37 treated bacteria showed compacted nucleoids and evidence for the formation of dense cytoplasmic structures (**Fig. 4E**). Because DNA and ribosomes are the main components of the bacterial nucleoid and cytoplasm, respectively, we tested whether LL-37 could condense these biomolecules in the test tube; it did (**Fig. 4F**). Thus, we suggest that the intracellular condensation caused by LL-37 is responsible for its bactericidal activity. Since LL-37 condenses with negatively charged biological structures, similar to PR, this condensation can be described according to the framework of so-called complex coacervation—originally conceptualized by the polymer physics field to describe the condensation of oppositely charged (bio)polymers^74,75^.

**Figure 4:**
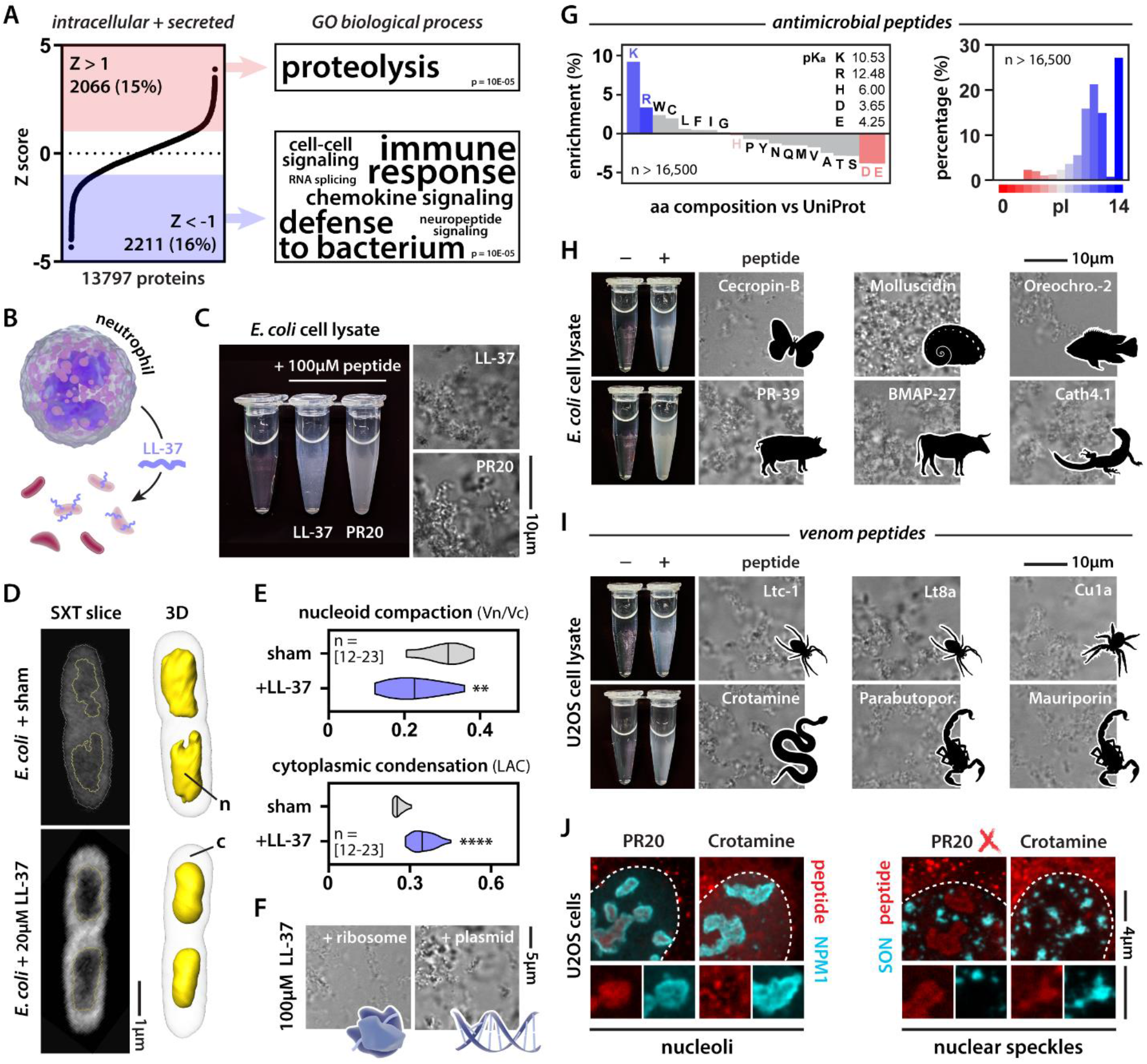
Killer peptides drive coacervation of cellular components and target biomolecular condensates. (A) The intracellular + secreted proteome ranked according to normalized confidence scores (Z score). GO categories associated with predicted red and blue proteins. (B) Scheme illustrating LL-37 as an AMP. (C) PR20 and LL-37 induce condensation of *E. coli* lysate. (D) Soft x-ray tomography of sham and LL-37-treated *E. coli* cells. n = nucleoid, c = cytoplasm. (E) Quantification of nucleoid compaction and cytoplasmic condensation. Mann-Whitney. ** p-value < 0.01, **** p-value < 0.0001. See also **Fig. S6**. (F) LL-37 condenses both *E. coli* ribosomes and plasmid DNA. (G) Positive charge is a common feature among AMPs. (H-I) Examples of natural AMPs and venom peptides that induce cell lysate condensation. (J) Crotamine and PR30 (500 nM) target endogenous biomolecular condensates in human cells.

Is LL-37 a sole example of such a mechanism, or could this be more widespread? We analyzed thousands of curated AMPs^76^, revealing that positive charge is a common feature among them (**Fig. 4G**). We tested a set of cationic AMPs from various vertebrates and invertebrates and found that several induce condensation in our *ex vivo* assay (**Fig. 4H**). These results support our emerging hypothesis that positively charged peptides could offer broad-spectrum antibiotic functionality through aberrant condensation of essential cellular components.

While AMPs target microbes, nature has also evolved biochemical tools to target large multicellular organisms in the form of venoms. Although several well-known (paralytic) venom peptides have a clear mode of action—usually specifically binding and modulating ion channels—venoms are complex mixtures containing a range of other peptides. Among them, so-called necrotic peptides are cationic and seem mainly responsible for tissue damage and cell death^77^. Indeed, we found that cationic peptides from the venoms of scorpions, spiders, and rattlesnakes drive condensation in our *ex vivo* lysate assay (**Fig. 4I**). These results suggest that they have the biophysical capacity to condense, but do they exhibit this behavior in cells? Crotamine is a well-studied cell-penetrant rattlesnake peptide whose mode of action is incompletely resolved^78^. We added fluorescently-labeled crotamine to the medium of human cells (**Fig. 4J**). As a control, we added fluorescently labeled PR, which has been documented to traffic to nucleoli^55,79^. PR targeted the granular center of the nucleolus, consistent with other reports^28,56^, while crotamine targeted nucleolar subcompartments. PR did not target nuclear speckles but crotamine did. Both these potent killer peptides—one aberrantly produced from an expanded repeat in disease, one evolved in a venomous animal—similarly targeted RNA-rich nuclear bodies. Yet despite these similarities, distinct structural and sequence features determined condensate partitioning, providing evidence that condensate target specificity can be encoded in the sequences of even small peptides.

### Cationic charge is necessary and sufficient for killer activity

Both the arginine-rich repeat peptides in human disease and the natural killer peptides have one main feature in common: positive charge. Could it be that charge is necessary and sufficient for killer activity? To address the first question, we looked more deeply at the specific amino acid enrichments we had seen above. While AMPs are enriched for arginine- and lysine-rich sequences, we did not see any enrichment for histidine (**Fig. 4G**)^76^. This observation is not unexpected, given histidine is expected to be largely neutral at a pH of 7.0 – 7.4. Nonetheless, two of our top ten secreted predicted blue proteins belonged to the histatin family—a histidine-rich group of AMPs (**Fig. S5B**). When tested in the same assay as LL-37, histatin-3 (HTN3) did not induce condensation in our *ex vivo* assay. However, histatins are expressed specifically in saliva, which is slightly acidic (≍ pH 6.5). Switching assay conditions from pH 7.5 to pH 6.5—coinciding with a net charge gain of +3—drove the strong condensation of HTN3 in *E. coli* lysate. This finding provides evidence that cationic charge is required for condensation, and is consistent with the enhanced bactericidal activity of histatins under mildly acidic conditions first observed three decades ago^80^. Thus, evolution seems to have used histidine residues to generate pH-dependent condensation switches that tune the activity of AMPs to their respective biofluid of origin.

Our work thus far suggests a net positive charge is necessary for toxicity of numerous AMPs, but is it sufficient? In other words, can we make a non-toxic protein toxic simply by altering its charge? GFP is probably the best-studied protein, with predictable behavior and folding, and is generally non-toxic to cells. By mutating residues facing out from the β-barrel, one can generate GFP mutants that still fold properly (i.e., remain fluorescent) but have an altered surface charge (**Fig. 5A**)^81^. When we expressed a library of super folder GFP (sfGFP) variants—with net charges ranging from -24 to +24—in *E. coli*, we observed that cationic GFP variants caused bacterial growth arrest (**Fig. 5B**). Imaging these cells, we observed that the cationic GFP variants, but not the anionic ones, formed aberrant condensates with ribosomal proteins (**Fig. 5C**) and RNA^82^ at the bacterial cell pole. Similarly, when we combined recombinantly purified GFP variants with *E. coli* ribosomes in the test tube, only the cationic ones formed condensates (**Fig. 5D**). Thus, like we saw for LL-37, these GFP variants can drive charge-based coacervation of ribosomes—killing the cells.

**Figure 5:**
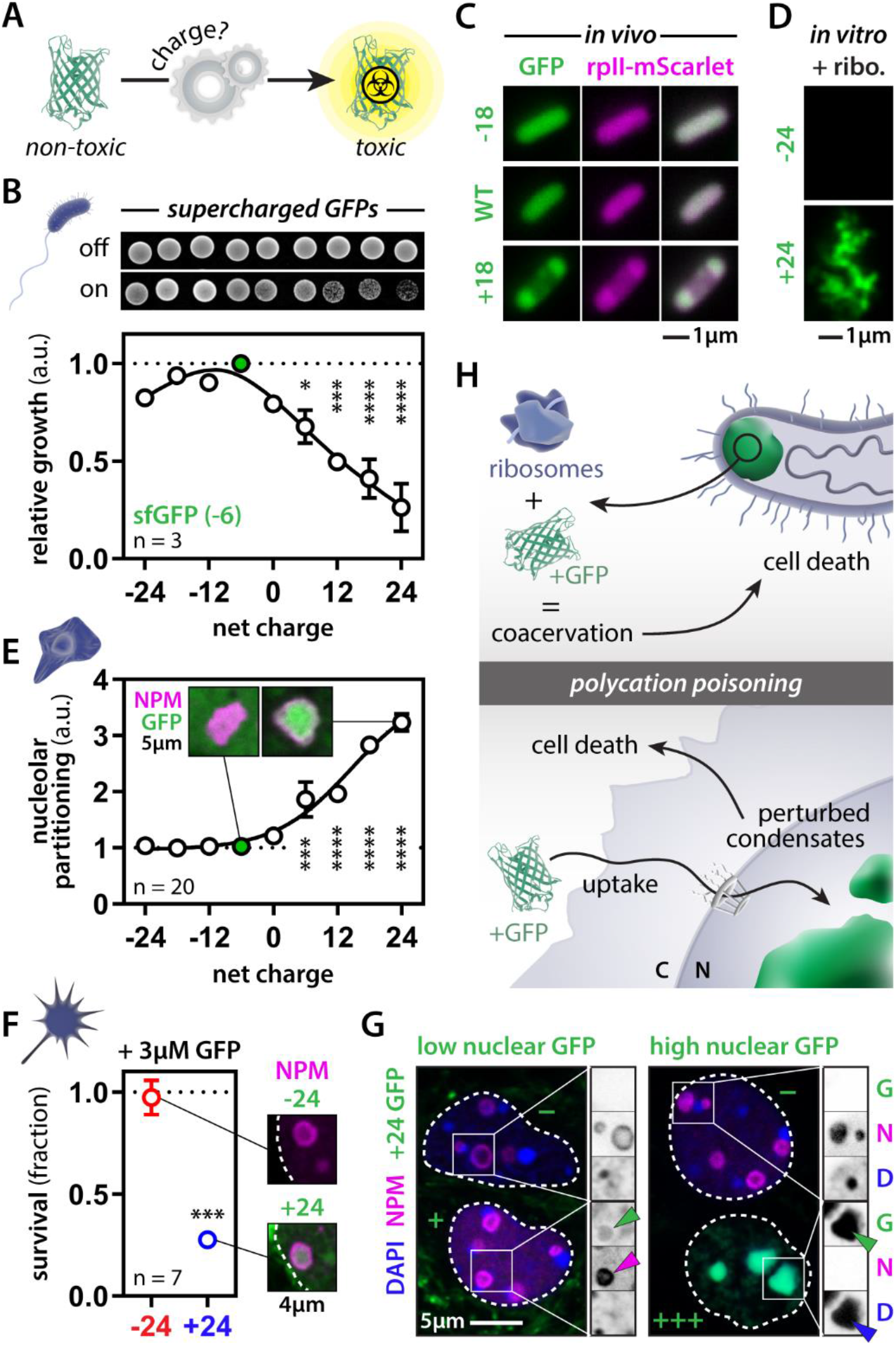
Net positive charge is sufficient for protein toxicity. (A) Engineering supercharged GFP versions for toxicity assays. (B) Expression of cationic GFP variants drives toxicity in *E. coli*. One-way ANOVA. (C-D) Cationic GFPs condenses with ribosomal proteins in *E. coli* and the test tube. (E) GFP partitions into nucleoli of U2OS cells in a positive charge-dependent manner. One-way ANOVA. (F) Exogenous recombinant cationic GFP is taken up, targets nucleoli and kills mouse cortical neurons in the dish. Student’s t-test. (G) Cells with low levels of GFP (+) have a normal nuclear organization, whereas high levels (+++) perturb nucleoli and heterochromatin. Adjacent cells that did not take up GFP (-) are normal. (H) Scheme depicting polycation poisoning in pro- and eukaryote systems. * p-value < 0.05, *** p-value < 0.001, **** p-value < 0.0001.

We next tested the effect of these GFP variants in human cells. The cationic GFP variants spontaneously condensed in nucleoli and nuclear speckles in a charge-dependent fashion (**Fig. 5E**). Given cationic peptides often undergo spontaneous cellular entry, we wondered if our synthetic positively charged GFPs might also be sufficient for cellular uptake. To test this, we also added recombinant anionic and cationic GFP to the medium of cultured mouse cortical neurons. Virtually identical to our observations with PR and crotamine, cationic GFP was spontaneously taken up by cells and targeted the nucleolus, killing the neurons (**Fig. 5F**). Moreover, this cationic version of GFP did so at concentrations similar to or lower than the ones we and others have found GR to be toxic^49,55,79^. Importantly, this observation challenges that canonical expectation that cell-penetrant peptides must have evolved a specific and highly constrained mechanism for cellular uptake, instead implicating cellular entry as being driven in part if not entirely by a strong net positive charge. Finally, neuronal uptake of cationic GFP was not uniform; some cells rapidly accumulated large quantities of GFP, whereas in others the accumulation was slower or even absent. Note, this same heterogeneity has been observed for other cell-penetrant peptides as well^83–85^ and is linked to cellular tolerance against their toxic effects^83^. Cells with low levels of nuclear GFP showed nucleolar localization as expected (**Fig. 5G**). Surprisingly, cells with high levels of GFP had non-detectable nucleoli (based on NPM staining) and showed marked condensation of DNA. The oversaturation of the nuclear compartment with these charged molecules seems to upset the physiological balance of electrostatic interactions, dramatically perturbing nuclear organization and leading to the demise of the cell (**Fig. 5H**).

Taken together, our results suggest that positively charged peptides and proteins offer broad-scale cellular toxicity. In some cases, this toxicity is an evolved function, while in others, it may be an unfortunate mishap through spontaneous repeat expansions that underlie human disease. Our results demonstrate that while precise target selectivity can clearly be encoded in a killer peptide’s sequence (e.g., crotamine vs PR), we conclude that cationic charge is a unifying sequence feature that is sufficient to drive killer activity. This process, which we name *polycation poisoning*, follows the simple framework of complex coacervation and has independently evolved countless times across the tree of life.

## DISCUSSION

For a decade, the observed target promiscuity of C9orf72 arginine-rich DPRs has puzzled the field. Compelling data has connected their toxicity to a bafflingly diverse array of biological pathways^20–22^. But a unifying framework has so far remained elusive. Were these studies monitoring the cause of cell death or simply the downstream consequences of dying cells? We and others had previously shown that these peptides can condense RNA and other biomolecules, explaining their effect on physiological condensates^22,27,28,33^. Here, we used proteomics to explore the target space of cationic dipeptide repeats and unexpectedly found that even their non-condensate targets fit into a simple electrostatics-based framework of complex coacervation. We propose that key actors in all these different biological pathways share the right sequence features that drive their aberrant interaction with these positively charged peptides.

Our framework also allows us to address the most common criticism against the pathological role for arginine-rich DPRs in patients. In every model system tested so far, PR—and to a lesser extent GR— targets the nucleolus, making it potently toxic. Yet, this has never been observed in any pathology report using patient post-mortem material. For reasons that are currently unresolved, but a topic of intense study, both PR and GR seem to get trapped in the cytoplasmic GA inclusions in patients, thus preventing them from entering the nucleus^17–19^. Our results unambiguously demonstrate that RNA tunes the target space of PR (**Fig. 1**). As such, which proteins PR (and, by extension GR) interact with within the cell will depend on the local RNA concentration around those DPRs. The concentration of RNA in the nucleus is estimated to be 36 times higher than the cytoplasm^44^, where most RNA is also trapped in polysomes. As such, our test tube experiments provide lysate mimics of either a nucleus-like (high RNA) or cytoplasm-like (low RNA) environment, enabling us to dissect which proteins are PR targets in these two compartments. Under high RNA conditions, we observe PR interacting with proteins associated with nuclear bodies and chromatin. This mirrors the subcellular localization observed in model systems, where PR interacts with nucleoli^27,28,55,56^ and heterochromatin^57^. Under low RNA concentrations, PR interacts with the translation initiation complex, ribosomes, axonal transport factors, and protein arginine methyl transferases. Analysis of patient post-mortem samples provides evidence that these kinds of proteins do associate with cytoplasmic GR/PR inclusions in patients^24,30,49,52,53^. Although defects in condensation seemed initially limited to disease models that failed to fully recapitulate human pathology, we now show that the same framework of electrostatic condensation is still applicable *but* is tuned by the local RNA concentration. To our knowledge, this is the first model that is able to explain a decade of disconnect between most C9orf72 disease model-based studies and human pathology reports.

Pathogenesis does not exist in a vacuum, but instead follows already-established biochemical principles. Can we use a framework designed to understand pathology to infer novel functional biology? We used our proteomics dataset to train a machine learning algorithm to detect similar electrostatics-based condensation behavior. When tested on the human proteome, we found already known condensate proteins but, importantly, also predicted this behavior in completely uncharacterized proteins. We experimentally validated this condensate behavior in nine out of the ten predicted proteins—showing that our approach allows for the discovery of new biology. The complete functional characterization of this set of novel condensate proteins is beyond the scope of this study, but we note that some of these proteins have substantial phenotypic effects in high-throughput knock-out studies^86^, arguing that their yet unknown function may carry cellular importance. It will be interesting to test if their condensation propensity is involved in their function, and whether they are implicated in human disease.

The (re)discovery of biomolecular condensation as an organizing principle of the cell besides membranes has revolutionized cell biology^87,88^. Ironically though, our exploitation of these insights has typically remained delimited by the plasma membrane. With our sequence-based condensate protein predictor in hand, we could now test for the condensation properties of extracellular proteins in a direct and unbiased way. To our initial surprise, we discovered that cationic killer peptides found in innate immune systems and venom cocktails are potent drivers of condensation, following the exact same set of rules we delineated for the human disease repeat peptides. Not impeded by GA inclusions like PR, venom peptides are able to gain entry to mammalian cells and readily target the nuclear centers of RNA production. AMPs condense both the nucleoid and ribosomes in bacteria. Our results demonstrate that two very different cellular systems (eukaryotes versus prokaryotes) with completely unrelated peptides follow the exact same framework of polycation poisoning. Indeed, using GFP as a scaffold we can engineer killer activity by simply making it positively charged, with its toxicity in neuronal culture rivaling that of the neurotoxic GR and PR peptides.

One question remains though: why positive and not negative charge? Although we do not rule out the existence of polyanion-based toxicity, there are some simple reasons why cationic peptides could be more disruptive. If we consider the cell as a collection of lipids, nucleic acids and proteins, a clear trend emerges. Lipids tend to have negatively charged head domains, nucleic acids are inherently anionic, but proteins can span the entire charge spectrum. Given all the negative charge on lipids, nucleic acids, and even the cytoskeleton (both actin filaments and microtubules have negatively charged surfaces), a cationic-biased proteome would promiscuously interact with all of these exposed biomolecular surfaces. Hence, it may come as no surprise that for organisms living at mesophilic pH and salt concentrations, proteomes tend to be—on average—more negatively charged^89,90^. Yet, we know many examples of highly cationic proteins in the cell, raising the question of how cells protect themselves from their own. By analyzing six model organisms, we indeed find no evidence for selection against positively charged proteins (**Supplemental text**, **Fig. S7**), despite our data suggesting that polycationic folded and disordered proteins drive cellular toxicity across the kingdoms of life. To better understand these seemingly contradictory observations, we performed an integrative bioinformatic analysis showing that while proteomes do possess many examples of proteins with a net positive charge, these are either expressed at low copy numbers, sequestered from the cellular environment via constitutive binding to anionic biomolecules, or neutralized through posttranslational modifications (**Supplemental text**, **Fig. S8-S11**, **Table S3**). Remarkably, we find that for essentially every single example of an abundant, positively charged protein, its charge is neutralized by a phosphate moiety—either in *trans* (nucleic acids or phospholipid headgroups) or in *cis* (through phosphorylation). We argue that this neutralization is essential to keep endogenous polycationic proteins in check to prevent them from poisoning the cell. Intriguingly, similar approaches are used by several human pathogens to neutralize cationic antimicrobial peptides and antibiotics^91–93^.

The above observations point at a seemingly universal proteome ‘design principle’. The net polarity for the large cellular surfaces ensures cellular proteomes do not aspecifically stick to lipids and nucleic acids. At the same time, it becomes relatively easy to evolve high-affinity domains that engage with these acidic moieties if needed. Cationic surface patches or disordered domains are pervasive interaction motifs in the proteome. For example, countless RNA binding proteins possess cationic disordered domains adjacent to their folded RNA binding domains^27,94,95^. Recent work has shown that even modest clusters of positively charged residues in IDRs from RNA binding proteins can have a profound impact on RNA binding affinity, mirroring our bioinformatic analyses^96^. Similarly, positively charged microtubule-binding proteins, such as tau, do so in part by interacting with the anionic tubulin tails that coat the microtubule lattice^97^. Membrane-binding proteins, like α-synuclein, do so via the exact same electrostatic interactions^98^. The use of cationic interaction and targeting motifs in a sea of anionic biomolecules is a powerful one but highlights the inherent weakness of cells against killer systems that corrupt this principle. It also shows why the cell has a variety of fail-safe mechanisms in place to prevent toxicity of its own cationic proteins. When we tested the cationic disordered domains of some human nucleic acid binding proteins in isolation, we found they act virtually identical to the killer peptides described in this study (**Fig. S12**). So how does the cell tame these sequences? Above we highlight the role that phosphorylation could have in regulating these proteins. Recent studies by us and others have also shown that these cationic domains are often paired with anionic or aromatic domains that can potently quench or regulate their activity^99–103^. Additionally, nuclear import factors that bind proteins translated in the cytoplasm via their nuclear localization sequence—usually cationic in nature—were recently shown to moonlight as *bona fide* protein chaperones^104–107^. Although philosophical, it is worth considering which function came first when transcription and translation became uncoupled during eukaryogenesis: chaperoning a condensation-prone protein in an RNA-poor environment (i.e., the cytosol) or translocating it to an RNA-rich environment (i.e., the nucleus)?

In this study, we provide a simple electrostatics-based framework that unifies the toxicity of cationic peptides in human disease and killer peptides in biology. McKnight and co-workers speculated that RAN translation of the *C9orf72* repeat could be considered the “failed birth of a gene”^55^. Upon repeat expansion, a non-coding part of the genome all of a sudden becomes protein-coding. Yet, the gene product is toxic to the cell producing it. When venomous animals evolved their killer peptides, a requirement was that the venom did not kill its host. Unsurprisingly, venom peptide production is a tightly regulated process often involving extracellular processing and cleavage of quenching domains together with signal peptides correctly targeting the protein for secretion. Note that in the lumen of a secretory vesicle these peptides never “see” the cytoplasm of the cell they are produced in. With this in mind, one can only wonder whether we have witnessed the failed birth of a venom peptide that lacked its regulatory sequences.

## ACKNOWLEDGEMENTS

We thank all members of the Gitler lab, the Boeynaems lab, the Carnegie-Stanford Intrinsically Disordered Protein Scientific Interest Group (IDPSIG), IDPseminars, and FNZ for helpful discussion and suggestions. We are indebted to Dr. Bede Portz for providing invaluable feedback and insights that helped shape this manuscript. We thank the Stanford Neuroscience Microscopy Service for use of the core facility. Some of the computing for this project was performed on the Sherlock cluster. We would like to thank Stanford University and the Stanford Research Computing Center for providing computational resources and support that contributed to these research results. We thank members of the Water and Life Interface Institute (WALII), supported by NSF DBI grant # 2213983, for helpful discussions. We thank Dr. Richard Kriwacki and Dr. Aaron Phillips (St. Jude Children’s Research Hospital) for kindly providing us with recombinant NPM1.

## AUTHOR CONTRIBUTIONS

**S.B:** Conceptualization, Methodology, Validation, Formal Analysis, Investigation, Data Curation, Writing-Original Draft, Visualization. **X.R.M.:** Software, Formal Analysis. **V.Y.:** Investigation, Formal Analysis. **G.G.:** Software, Formal Analysis. **JH.C.:** Investigation. **J.B.:** Software, Formal Analysis. **L.N.:** Investigation. **A.S.:** Investigation, Formal Analysis. **A.B.:** Investigation. **D.VH.:** Investigation. **J.P.:** Investigation. **A.E.:** Formal Analysis. **H.B.S:** Formal analysis. **K.S.:** Resources. **L.P.:** Methodology. **K.L.:** Methodology, Formal analysis. **C.L.:** Supervision. **M.A.F.H.:** Methodology. **A.K.:** Supervision. **F.I.:** Methodology, Supervision. **A.O.:** Methodology, Supervision. **A.S.H.:** Methodology, Formal analysis, Supervision, Writing-Review & Editing. **A.D.G.:** Conceptualization, Supervision, Funding acquisition, Writing-Review & Editing.

## MATERIAL & METHODS

### EXPERIMENTAL MODELS

#### Human cell lines

U2OS (ATCC, HTB-96) cells were grown at 37°C in a humidified atmosphere with 5% CO_2_ for 24 h in Dulbecco’s Modified Eagle’s Medium (DMEM), high glucose, GlutaMAX + 10 % Fetal Bovine Serum (FBS) and pen/strep (Thermo Fisher Scientific).

#### Bacteria

*E. coli* (Stbl3, Thermo Fisher) were grown at 37°C in a shaking incubator in LB medium and used for ex vivo and soft x-ray tomography assays. *E. coli* (NiCo21(DE3), New England Biolabs) transformed with plasmids for expression of GFP charge variants were grown at 37 °C in a shaking incubator in LB medium supplemented with 100 µg/mL ampicillin.

#### Mice and primary mouse cells

All mouse husbandry and procedures were performed in accordance with institutional guidelines and approved by the Stanford Administrative Panel on Animal Care (APLAC). C57BL/6 mice (Jackson Laboratory) were used. Mouse primary cortical neurons were grown in Neurobasal media (Gibco) supplemented with B-27 serum-free supplement (Gibco), GlutaMAX, and Penicillin-Streptomycin (Gibco) in a humidified incubator at 37°C, with 5% CO_2_.

### METHOD DETAILS

#### Plasmid construction

All constructs for human expression were generated through custom synthesis and subcloned into a pcDNA3.1 backbone by Genscript (Piscataway, USA). All constructs for GFP expression in *E. coli* were generated through custom synthesis and subcloned into an expression backbone (pETDuet-1) by Genscript (Piscataway, USA), as has been previously reported^110^.

#### Peptides

All peptides were generated via chemical synthesis by Pepscan (Lelystad, the Netherlands), except BMAP-27 (AS-65598, Anaspec), LL-37 (AS-61302, Anaspec), 5-FAM-LC-LL-37 (AS-63694, Anaspec), Crotamine (CRO01-01000, Smartox-Biotech), and Cy3-Crotamine (CRO02-00500, Smartox-Biotech). Peptides were fluorescently labeled as described previously ^111^.

#### Recombinant proteins

##### Recombinant GFP variants

GFP variants were expressed in *E. coli* (NiCo21(DE3), NEB) in LB media supplemented with 100 µg/mL ampicillin. Overnight cultures were inoculated into 1 L of LB media supplemented with ampicillin and allowed to grow at 37 °C to an OD_600_ between 0.8-1.0. At this point, the cultures were induced with 1 mM IPTG. After induction, cultures were grown at 25 °C with shaking for an additional 16-18 h. At this point, cells were collected by centrifugation in a swinging bucket rotor at 4000 rpm for 15 min. The cell pellet was resuspended in a lysis buffer (50 mM NaH_2_PO_4_, pH 8.0 supplemented with 300 mM (GFP(-24)) or 1 M NaCl (GFP(+24))). The cell pellet was then lysed by sonication (2s on, 4 s off) at 60% amplitude using a 0.5 in probe for 10 min. The insoluble fraction was separated via centrifugation in a fixed angle rotor at 10,000 rpm for 30 min at 4 °C. The GFP was purified from the soluble fraction using immobilized metal affinity chromatography (His-Pur Ni-NTA). The column was washed with lysis buffer containing 50 mM imidazole and GFP was eluted with lysis buffer supplemented with 250 mM imidazole. Fractions were collected and analyzed by SDS-PAGE for purity. Pure fractions were combined and dialyzed against 10 mM tris buffer, pH 7.4 before adjusting the concentration to 1 mg/mL GFP by ultrafiltration using a 10 kDa molecular weight cutoff spin filter (Amicon).

##### *In vitro* chromatin reconstitution

Chromatin was reconstituted by salt dialysis as described previously^112^. Histones for chromatin assembly were purified as described previously^112–114^. pUC19 plasmid DNA containing 19x “601” Widom positioning sequence^115^ (ASP 696) was digested with EcoRI, XbaI, DraI and HaeII (New England Biolabs). 19x 601 array was then purified using polyethylene glycol (PEG) precipitation followed by dialysis into TE buffer (10 mM Tris pH 8.0, 0.5 mM EDTA). EcoRI and XbaI 5’ overhangs were filled in with dGTP (New England Biolabs), dCTP (New England Biolabs), dTTP (New England Biolabs), and biotin-14-dATP (Thermo Fisher Scientific) using Large Klenow fragment 3’-5’ exo- (New England Biolabs). Biotinylated 19x 601 array DNA, H2A/H2B histone dimer, and H3/H4 tetramer were added to high salt buffer (10 mM Tris-HCl, pH 7.5; 0.25 mM EDTA; 2 M NaCl) and gradually dialyzed over the course of ∼67 hours at a rate of 0.5 mL/min from 500 mL high salt buffer into low salt buffer (10 mM Tris-HCl, pH 7.5; 0.25 mM EDTA; 2.5 mM NaCl). H3/H4 tetramer concentrations were titrated to vary chromatin saturation. Nucleosome assembly was verified using overnight digestion of chromatin at room temperature using AvaI which cuts between adjacent 601 positions in the array, followed by a native acrylamide gel shift analysis (gel shift of ∼200 bp AvaI digestion product to ∼600 bp suggests nucleosome positioned on 601 sequences as described previously^112^).

##### Recombinant NPM1

Recombinant NPM was a kind gift from Dr. Richard Kriwacki and Dr. Aaron Phillips (St. Jude Children’s Research Hospital).

#### Human cell culture, treatments and microscopy

U2OS cells were grown at 37°C in a humidified atmosphere with 5 % CO2 for 24 h in Dulbecco’s Modified Eagle’s Medium (DMEM), high glucose, GlutaMAX + 10 % Fetal Bovine Serum (FBS) and pen/strep (Thermo Fisher Scientific). Cells were transiently transfected using Lipofectamine 3000 (Thermo Fisher Scientific) according to manufacturer’s instructions. Cells grown on cover slips were fixed for 24 h after transfection in 4 % formaldehyde in PBS. For peptide uptake experiments, cells were treated with 1 µM of fluorescently labeled peptides and incubated for 45 min at 37°C, followed by a PBS wash and fixation with 4 % formaldehyde in PBS. Slides were mounted using ProLong Gold antifade reagent (Life Technologies). Confocal images were obtained using a Zeiss LSM 780 Meta NLO confocal microscope. Images were processed and analyzed using Fiji^116^ and Adobe Photoshop.

#### Mouse primary neuron culture and microscopy

Primary mouse cortical neurons were dissociated into single cell suspensions from E16.5 C57BL/6 mice (Jackson Laboratory) cortices using a papain dissociation system (Worthington Biochemical Corporation). Neurons were seeded onto poly-L-lysine coated plates (0.1% w/v) and grown in Neurobasal media (Gibco) supplemented with B-27 serum-free supplement (Gibco), GlutaMAX, and Penicillin-Streptomycin (Gibco) in a humidified incubator at 37°C, with 5% CO2. Cells were transiently transfected using Lipofectamine 3000 (Thermo Fisher Scientific) according to manufacturer’s instructions. Cells grown on cover slips were fixed for 24 h after transfection in 4 % formaldehyde in PBS. For peptide uptake experiments, cells were treated with 1 µM of fluorescently labeled peptides and incubated for 1 h at 37°C, followed by a PBS wash and fixation with 4 % formaldehyde in PBS. Slides were mounted using ProLong Gold antifade reagent (Life Technologies). Confocal images were obtained using a Zeiss LSM 780 Meta NLO confocal microscope. Images were processed and analyzed using Fiji^116^ and Adobe Photoshop.

#### Bacterial culture and microscopy

*E. coli* (NiCo21(DE3), New England Biolabs) transformed with plasmids for expression of GFP charge variants were grown at 37 °C in a shaking incubator in LB medium supplemented with 100 µg/mL ampicillin. For each variant, overnight cultures were prepared by inoculating a single colony from a plate into a snapcap tube containing 5 mL of LB media supplemented with ampicillin. After growth overnight, the cultures were then diluted to OD_600_ = 0.1 in LB media supplemented with ampicillin and were allowed to grow at 37 °C with shaking until they reached an OD_600_ of approximately 1.0. The OD_600_ of each culture was then normalized to 0.9 and 200 µL of the culture was then transferred to a 96-well plate for 10-fold serial dilutions with LB media supplemented with ampicillin. 5 µL of each culture and dilution (10-10^7^x diluted) were spotted onto LB plates supplemented with 100 µg/mL ampicillin with or without 1 mM isopropyl β-D-1-thiogalacopyranoside (IPTG, Gold Biotechnology). After drying, plates were incubated at 25 °C for 30 h before imaging on a scanner and transilluminator (Bio-Rad Gel Doc XR+).

For imaging GFP co-localization with ribosomes in *E. coli*, an rplI-mScarlet fusion was integrated into the native genome locus of rplI in the NiCo21(DE3) *E. coli* strain. This process began with QC101 (MG1655 *rplI- mCherryKan*R) cells, which contain a genomically integrated ribosome-mCherry fusion protein and were previously used to investigate the localization of ribosomes in *E. coli* during various phases of cell growth^117^. We received these cells as a gift from the Sanyal lab and amplified the rplI-mCherry_KanR cassette via colony PCR using primers, CGAACACGAAGTGAGCTTCC and AAGCAAAACGCCGACCAA, and Phusion polymerase as recommended by New England Biolabs (NEB). mCherry was replaced with mScarletI for better compatibility with our imaging system. HiFi assembly was used to assemble a vector with a rplI-mScarletI_KanR cassette.

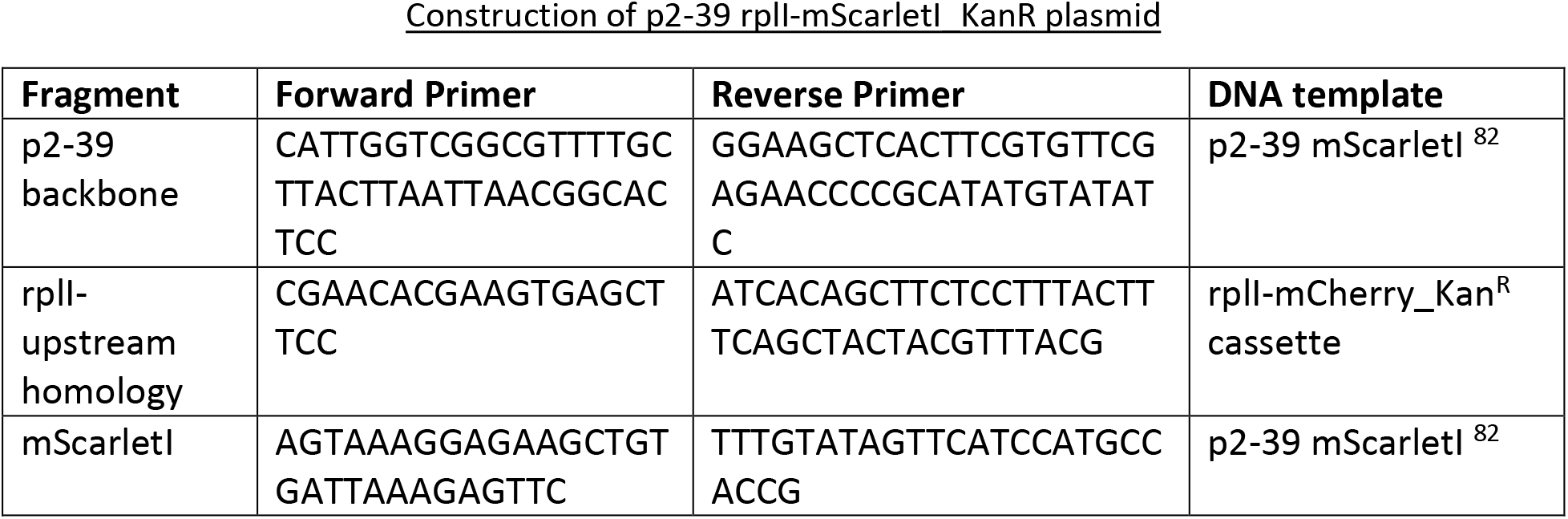
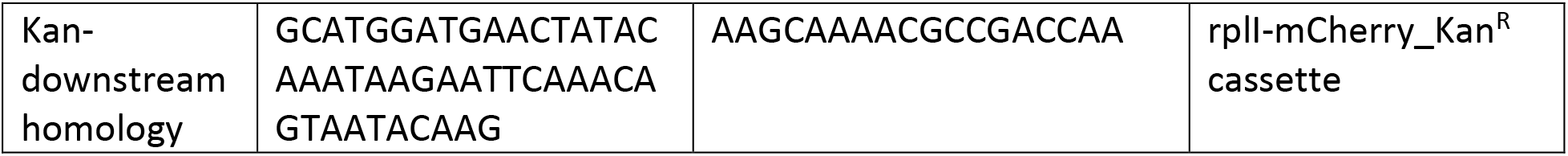

All fragments were PCR purified (NEB), ligated using NEB HiFi assembly kit, transformed into NEB5α cells according to the manufacturer’s instructions, and plated onto LB agar plates supplemented with 25 µg/mL chloramphenicol and 50 µg/mL kanamycin. Single colonies were inoculated into LB media supplemented with chloramphenicol and kanamycin and grown at 37 °C overnight with shaking. Plasmid DNA was isolated from overnight cultures using the Qiagen Spin Miniprep Kit and plasmid sequences were verified by Genewiz.

The dsDNA cassette for genomic integration was amplified from the p2-39 rplI-mScarletI_KanR plasmid using primers, CGAACACGAAGTGAGCTTCC and AAGCAAAACGCCGACCAA. The PCR-amplified product was treated with DpnI, PCR purified, and verified by gel electrophoresis.

To prepare cells for genomic integration, electrocompetent NiCo21(DE3) cells (NEB) were prepared and transformed with 50 ng pTKRED. pTKRED was a gift from Edward Cox and Thomas Kuhlman (Addgene plasmid # 41062)^118^. Briefly, overnight cultures of NiCo21(DE3) cells were diluted 100-fold in LB media and grown at 37 °C, 225 rpm until OD_600_ = 0.35-0.4. Cells were then incubated on ice for 10-20 min and centrifuged at 400 rcf for 10 min at 4 °C. Cells were washed twice in ice cold sterile milliQ water and then washed once in ice cold 10% glycerol before resuspending in ice cold 10% glycerol, aliquoting, and snap freezing with liquid nitrogen.

Electrocompetent NiCo21(DE3) cells were transformed with pTKRED using a MicroPulser Electroporator (Bio-Rad). 1 µL plasmid was added to 100 µL electrocompetent cells in a chilled Gene Pulser/MicroPulser electroporation cuvette with 0.1 cm gap (Bio-Rad) and incubated on ice for 1 min. Cells were then subjected to a single pulse and were then recovered in SOC media at 30 °C with shaking for 3 h. A portion of the recovered cells was then plated on LB plates supplemented with 50 µg/mL spectinomycin and incubated at 30 °C overnight. Single colonies were inoculated into LB media supplemented with 100 µg/mL spectinomycin and the presence of the plasmid was confirmed by sequencing (Genewiz).

Electrocompetent pTKRED NiCo21(DE3) cells were prepared. Cells were grown in SOB media supplemented with 0.5% glucose, 100 µg/mL spectinomycin, and 2 mM IPTG at 30 °C. Cells were then harvested as described above when OD_600_ = 0.5 – 0.6. 1 µg of the dsDNA rplI-mScarletI-KanR cassette was added to 100 µL electrocompetent cells, and electroporation was performed as described above. Cells were recovered in 1 mL SOC media for 3 h at 30 °C with shaking and 250 µL of electroporated cells were plated onto LB agar plates supplemented with 100 µg/mL spectinomycin and 25 µg/mL kanamycin^118^. Plates were incubated at 30 °C for 1-2 days. Single colonies were selected and grown in LB media supplemented with 25 µg/mL kanamycin and incubated at 42 °C with shaking for approximately 20 h before they were diluted and spread onto LB plates supplemented with 25 µg/mL kanamycin. Plates were incubated overnight at 37 °C. We verified that these colonies did not grow on LB plates with spectinomycin at 30 °C, indicating that the pTKRED plasmid had been cured. Additionally, we performed colony PCR with primers upstream (CGCCATCAGTAATCGGTCA) and downstream (CGCGAAGTTCTTCCACGAT) of the cassette, confirming successfully integration into the *E. coli* genome.

Chemically competent rplI-mScarletI_KanR NiCo21(DE3) cells were prepared and transformed with a plasmid encoding each GFP variant. Briefly, overnight cultures were diluted 100-fold and grown to OD_600_ = 0.3 -0.4 at 37 °C. At this point, cells were harvested by centrifugation at 2500 rcf for 10 min at 4 °C in a cold fixed angle rotor. The pellet was washed once in ice-cold transformation buffer (10 mM PIPES, 15 mM CaCl_2_, 250 mM KCl, 55 mM MnCl_2_, pH 6.7) and DMSO was added dropwise to a final concentration of 7%. Competent cells were aliquoted and snap frozen in liquid nitrogen. 2 µL of GFP plasmid (GFP(-18) or GFP(+18))^82^ was added to 100 µL chemically competent cells. Cells were incubated on ice for 30 min, heat shocked at 42 °C for 30 s, and incubated on ice for 5 min. Cells were recovered in 200 µL SOC for 1 h at 37°C and 100 µL of this mixture was plated onto LB plates supplemented with 50 µg/mL kanamycin and 100 µg/mL ampicillin. Cells were grown overnight at 37 °C.

For each variant, cultures were grown as above until they reached an OD_600_ between 0.8-1.0. At this point GFP expression was induced by the addition of 1 mM IPTG and the cultures were grown at 25 °C with shaking. Protein expression was induced for 24 h and then cells were imaged on agarose pads using a 100X oil 1.40 NA UPlanSApo objective (Olympus) on GFP (λ_ex_ = 470-522 nm; λ_em_ = 525-550 nm; EVOS GFP light cube) and Texas Red (λ_ex_ = 585-629 nm; λ_em_ = 628-632 nm; EVOS Texas Red light cube) channels using an EVOS FL Auto 2 inverted fluorescence microscope. Raw microscopy images were background subtracted (rolling ball algorithm with 100 pixel radius) in ImageJ.

#### Immunofluorescence

U2OS cells and primary neurons were fixed 24h after transfection (or after 45 min – 1 h of peptide treatment) in 4% formaldehyde in PBS and stained according standard protocols. Following primary antibodies were used: G3BP1 (ab56574, Abcam), H3K9me3 (ab8898, Abcam), SON (NBP1-88706, Novus Biologicals), FIBL (ab4566, Abcam), NPM1 (ab10530, Abcam), PABPC (ab21060, Abcam), FLAG (F7425 and F1804, Sigma-Aldrich). Alexa-labeled secondary antibodies were used (Thermo Fisher Scientific).

#### Neurotoxicity assays

Five days after seeding the neurons, recombinant GFP mutants were added at a concentration of 3 µM for 24h. Cytotoxicity in primary neuron cultures was measured with an alamarBlue cell viability (Thermo Fisher Scientific), according to manufacturer’s instructions. Readout was measured using a SPARK Multimode microplate reader (Tecan Life Sciences). Data was analyzed using Microsoft Excel and GraphPad Prism.

#### Peptide coacervation experiments

Peptide experiments were performed as described previously with slight modifications ^111^. In brief, peptides were generated via chemical synthesis by Pepscan (Lelystad, the Netherlands). Peptides were dissolved in milli-Q water and stored at -20°C. To test for coacervation, peptides were diluted in human or *E. coli* cell lysate at the indicated concentrations in PBS at pH 7.4. NPM1 phase separation assays were done as described in ^56^. Nucleosome arrays, plasmid DNA or E. coli ribosomes (New England Biolabs) were added at a concentration of 200 ng/µl and 1 µg/µl, respectively. Samples were transferred to an imaging chamber (Grace Bio-Labs, Bend, USA) and imaged on a Zeiss LSM 710 confocal microscope. Pictures of turbid solutions in PCR strips were taken using a Google Pixel 3a.

#### PR protein precipitation

U2OS cells were trypsinized (Thermo Fisher Scientific) and harvested by centrifugation. Cell pellets were washed three times with ice-cold PBS. Pellets were redissolved in PBS buffer with Halt Protease inhibitor (Thermo Fisher Scientific) and lysed using a probe sonicator (Branson Sonifier 250, VWR Scientific) on ice. The lysate was cleared from the insoluble fraction by centrifugation for 15 min at 10,000 rpm at 4°C. The supernatant was retrieved and the procedure was repeated until no pellet was visible after centrifugation. The protein concentration was measured using Micro BCA assay (Thermo Fisher Scientific). PR30 (Pepscan, The Netherlands) was added to a final concentration of 50 mM to 0.5 mg of soluble lysate in a total volume of 400 µl and incubated at room temperature for 15 min. The volume of the samples was increased to 1 ml with PBS, before gently spinning down the PR droplets at 4,000 rpm for 5 min. The supernatans was transferred to a new tube. Pellets were subsequently washed with 1 ml PBS and vortexed before spinning down again. Washing steps were repeated three times. To each pellet a volume of buffer corresponding to the supernatans was added. The resulting samples were processed for LC-MS/MS. Samples for analysis by silver staining were generated identically. Silver staining (Thermo Fisher Scientific) was performed according to the manufacturer’s instructions.

#### Proteomics sample preparation and LC-MS/MS analysis

Proteins in the supernatant and pellet samples were first reduced by addition of DTT to a concentration of 5 mM and incubation for 30 minutes at 55°C and then alkylated by addition of iodoacetamide to a concentration of 10 mM for 15 minutes at room temperature in the dark. Samples were diluted with 50 mM Tris pH 7.9 to a urea concentration of 4 M and proteins were digested with 5 µg lysyl endopeptidase (Wako) (1/100, w/w) for 4 hours at 37°C. Samples were further diluted with 50 mM Tris pH 7.9 pH 8.0 to a final urea concentration of 2 M and proteins were digested with 5 µg trypsin (Promega) (1/100, w/w) overnight at 37°C. Next, peptides were purified on SampliQ SPE C18 cartridges (Agilent). Columns were first washed with 1 ml 100% acetonitrile (ACN) and equilibrated with 3 ml of solvent A (0.1% TFA in water/ACN (98:2, v/v)) before samples were loaded on the column. After peptide binding, the column was washed again with 2 ml of solvent A and peptides were eluted twice with 750 µl elution buffer (0.1% TFA in water/ACN (40:60, v/v)). Purified peptides were dried under vacuum in HPLC inserts and stored at −20 °C until LC-MS/MS analysis.

Peptides were re-dissolved in 20 µl loading solvent A (0.1% TFA in water/acetonitrile (ACN) (98:2, v/v)) of which 2.5 µl (supernatant) or 5 µl (pellet) was injected for LC-MS/MS analysis on an Ultimate 3000 RSLC nanoLC in-line connected to a Q Exactive HF Biopharma mass spectrometer (Thermo Fisher Scientific) equipped with a pneu-Nimbus dual ion source (Phoenix S&T). Trapping was performed at 10 μl/min for 4 min in loading solvent A on a 20 mm trapping column (made in-house, 100 μm internal diameter (I.D.), 5 μm beads, C18 Reprosil-HD, Dr. Maisch) and the sample was loaded on a reverse-phase column (made in-house, 75 µm I.D. x 400 mm length, 3 µm beads C18 Reprosil-HD, Dr. Maisch). Peptides were loaded with loading solvent A and were separated with a non-linear 145 min gradient from 2% to 56% MS solvent B (0.1% FA in water/ACN 20:80 (v/v)) at a flow rate of 250 nl/min followed by a 15 min wash reaching 99% MS solvent B and re-equilibration with 98% MS solvent A (0.1% FA in water).

The mass spectrometer was operated in data-dependent mode, automatically switching between MS and MS/MS acquisition for the 16 most abundant ion peaks per MS spectrum. Full-scan MS spectra (375-1,500 m/z) were acquired at a resolution of 60,000 in the Orbitrap analyzer after accumulation to a target value of 3,000,000. The 16 most intense ions above a threshold value of 13,000 (minimum AGC of 1,000) were isolated for fragmentation at a normalized collision energy of 28%. The C-trap was filled at a target value of 100,000 for maximum 80 ms and the MS/MS spectra (200-2,000 m/z) were acquired at a resolution of 15,000 in the Orbitrap analyzer with a fixed first mass of 145 m/z. Only peptides with charge states ranging from +2 to +6 were included for fragmentation and the dynamic exclusion was set to 12 s. QCloud was used to control instrument longitudinal performance during the project^119^.

#### Protein identification and quantification

Data analysis was performed with MaxQuant (version 1.5.8.3) using the Andromeda search engine with default search settings including a false discovery rate set at 1% on PSM and protein level. The spectra of all LC-MS/MS runs were interrogated against the human proteins in the Swiss-Prot Proteome database (database release version of January 2018 containing 20,243 human protein sequences, (http://www.uniprot.org)). The mass tolerance for precursor and fragment ions was set to 4.5 and 20 ppm, respectively, during the main search. Enzyme specificity was set as C-terminal to arginine and lysine, also allowing cleavage at proline bonds with a maximum of two missed cleavages. Variable modifications were set to oxidation of methionine residues and acetylation of protein N-termini. Matching between runs was enabled with a matching time window of 0.7 minutes and an alignment time window of 20 minutes. Only proteins with at least one unique or razor peptide were retained leading to the identification of 1599 proteins. Proteins were quantified by the MaxLFQ algorithm integrated in the MaxQuant software. A minimum ratio count of two unique or razor peptides was required for quantification. The mass spectrometry proteomics data have been deposited to the ProteomeXchange Consortium via the PRIDE partner repository with the dataset identifier PXD040619.

#### Soft X-ray tomography

PR30 (Pepscan, The Netherlands) was added to a final concentration of 50 mM to 0.5 mg of soluble lysate in a total volume of 400 µl and incubated at room temperature for 15 min. Samples were subsequently loaded in thin-wall glass capillaries and rapidly frozen into liquid nitrogen cooled liquid propane. The specimens were imaged by XM-2, a soft x-ray microscope in the National Center for X-ray tomography (http://ncxt.org) located at the Advanced Light Source of Lawrence Berkeley National Laboratory (ALS, LBNL). The x-ray microscope operates at the “water-window” region of soft x-rays, providing natural contrast of biomolecules with respect to water. The condenser and objective lenses, chosen for this experiment, guaranteed isotropic 32nm voxel size^120,121^. For 3D reconstructions, 92 projection images, with 200-300 ms exposure time each, were acquired sequentially around a rotation axis with 2° increment angles. After normalization and alignment, tomographic reconstructions were calculated using iterative reconstruction methods^122^. The segmentation and visualization of multi-droplets was done based on linear attenuation coefficient of x-rays in Amira 6.3.0.

#### Clustering and machine learning

To group proteins according to their relative solubility profiles, we used both the three experimental solubility values and calculated the change in solubility between 0 and 0.4, and 0.4 and 2. These delta values were then clustered at several different k values using the kmeans algorithm. Aggregate solubilities were plotted and visualized, and k=9 was selected based on a high level of intra-cluster concordance and specificity of GO enrichment.

UniRep had been demonstrated to be an effective model to extract fundamental features of amino-acid sequences that are structurally, evolutionarily and biophysically grounded^64^. We obtained the UniRep average hidden state representations of the entire human proteome (UP000005640_9606) using the 256-unit model. We then set aside the proteins in the mass spec experiment to train a support vector machine model with a linear kernel (sklean.svn.SVC) that was able to distinguish blue from grey+red proteins. The mass spec proteins were randomly split into 80-20 train-validation sets. We then classified the rest of the proteome using the trained model, and the confidence scores were used to rank the proteins in each category.

Since we performed the mass spectrometry experiments on soluble cell lysate without the addition of detergents—to avoid any effects of the detergent on condensation—our input sample was strongly depleted from membrane proteins. Nonetheless, as can be seen in the GO analysis of cluster 9, we got a small membrane contaminant fraction specifically in the blue class of proteins. This makes sense as these membrane proteins were insoluble in all conditions of the experiment due to the lack of detergent, leading the algorithm to rank membrane proteins incorrectly with high blue scores. Therefore, we excluded all membrane bound proteins from our prediction analyses as our experimental setup was not designed to study the phase behavior of this class of proteins.

#### Charge clustering metric

The charge clustering metric implemented here was developed and adapted from an inverse weighted distance metric, which was originally developed in the context of clustering of aromatic residues^63^. This metric quantifies the clustering of target amino acids relative to other target residues, and is complimentary, yet distinct, from other current charge distribution metrics^59^. While other metrics quantify the relative patterning of charged residues, our focus here is not on the patterning of residues relative to charged residues being evenly distributed, but instead on identifying folded domains or disordered regions where positively or negatively charged residues are clustered together, regardless of the presence of well-mixed regions elsewhere in the same protein. The IWD metric also enables the same functional form to be used for folded domains and disordered regions. To adapt the clustering in the context of charge residues, a net-charge per residue (NCPR) is calculated for all charged residues using a sliding window of five residues. Next, positively charged or negatively charged residues are selected as so-called *target residues*. Then the NCPR of each target residue is weighted by the sum of the inverse distances between all pairs of other target residues for each residue (Fig. S3). To calculate a single value for the sequence an average across the target residues is computed. In the positive charge clustering, the target residues are the “true” positively charged (arginine and lysine) residues, where “true positive” here means the net charge of the residue is positive, as determined by the sliding-window NCPR. Conversely, in the negative charge clustering, the target residues are the “true” negatively charged residues (aspartic acid and glutamic acid) residues, where “true negative” again means the net charge of the residue is negative as determined by the sliding window NCPR.

To calculate the bivariate-charge clustering, a metric which reports on how well interspersed the positive and negative residues are relative to each other, the target residues are both true positive and true negative residues (again where ‘true’ here reflects the use of the sliding window NCPR). Clustering is then calculated only between pairs of opposite charge residues. The sum of the inverse distance between all pairs of oppositely charged target residues is then weighted by the absolute value of the difference in NCPR between target residues in the pair. In the IDRs, inverse distance is calculated in linear sequence space, while for the surface of folded domains inverse distance is calculated in cartesian space^123^.

#### Enrichment of protein sequence feature calculation

The protein sequence analysis was conducted using SHEPHARD, a Python-based framework designed for integrating and analyzing large-scale amino acid sequence properties^124^. IDRs were predicted and annotated using metapredict (V2)^125^. Protein structures for folded domains were taken from AlphaFold2 structure predictions^126^. The folded domain structures were analyzed to calculate the per-residue solvent-accessible surface area using SOURSOP and the 3D position of solvent-accessible residues were used to calculate 3D charge clustering (see above)^127^. Sparrow (https://github.com/idptools/sparrow) was used to calculate the sequence features for both IDRs and folded domains. While our analysis focused on charged residue clustering, we performed a full analysis of different sequence features to perform an unbiased screen to identify features that delineated grey, red, and blue class proteins. To determine which sequence features were most discriminatory in separating the three groups, an enrichment test was conducted comparing a given feature’s values to the whole dataset. To evaluate how well correlated a specific sequence feature is with a given class, a linear-discriminant analysis (LDA) was conducted. The weight coefficients from the LDA were then used to inform how well correlated a specific sequence feature was with a given group. This was conducted for both sequence features on the surface of the folded domains and in the IDRs.

#### Bioinformatics analysis of protein abundance

Protein abundance analyses (Fig. S7–11) were performed using SHEPHARD^124^ and sparrow (https://github.com/idptools/sparrow). Mass spectrometry data were obtained for humans^128^, *X. laevis*^129^, *A. thaliana*^130^, *E. coli*^131^, *S. pombe*^132^, and *S. cerevisiae*^133^. Phosphorylation data for the human proteome was obtained from ProteomeScout^134^. All data and code for this analysis are provided at https://github.com/holehouse-lab/supportingdata/tree/master/2023/boeynaems_2023.

#### Peptide charge calculation

Peptide charge as a function of pH was calculated using Protein Calculator v3.4 (http://protcalc.sourceforge.net/).

#### GO enrichment analysis

Go enrichment analysis was performed using DAVID^135^. Enriched terms are depicted in word clouds with font size correlated to -log(p-value). P-values were Bonferroni corrected.

### QUANTIFICATION AND STATISTICAL ANALYSIS

All data was analyzed using Graphpad Prism 8.4.1 and Excel. Statistical tests, p values, number of samples, replicates, and experiments are indicated in the figure legends.

## SUPPLEMENTAL TEXT

### Proteome-wide net charge analysis

We and others have previously indicated that proteomes tend to be—on average—more negatively charged^89,90,136^. We confirmed this conclusion in six different model organisms (**Fig. S7**). Since our work thus far suggests that polycationic folded and disordered proteins drive cellular toxicity across the kingdoms of life, one might expect a strong selection against positively charged proteins at the proteome level. To our surprise, bioinformatic analysis of these six different model organisms found little obvious evidence of this signature, with approximately equal numbers of positively and negatively charged proteins, despite the modest trend towards negatively charged proteins (**Fig. S7**). Yet, an implicit assumption in these analyses is that all proteins are equally abundant, which is obviously false. Using previously published quantitative mass-spectrometry data, we used the per-protein copy numbers to re- assess our analysis. Focusing on the set of proteins that comprise the top 85% of all proteins in a cell by abundance, we re-analyzed our proteomes using the overall biomolecular mass (i.e., copy number times protein molecular weight) (**Fig. S8**). We took this mass-weighted approach because we reasoned that poisons are intrinsically dose-dependent, such that we should expect an effect to depend on the overall mass of polycations, not simply the number of molecules. Our analysis identified 1063 highly abundant proteins, of which 154 (∼15%) had a net positive charge above +0.05. Of these polycationic proteins, 75% were histones or ribosomal proteins, 8% were membrane proteins, 6% were constitutive components of ribonucleoprotein bodies (e.g., the nucleolus, the spliceosome, or the signal recognition particle), and the remaining 11% were nuclear RNA- or zinc-binding proteins (**Fig. S9**, **Table S3**). A key feature of these proteins is that none exist as soluble cytosolic proteins; all but 24 are constitutively sequestered in assemblies with either nucleic acids (ribosomal proteins or histones) or phospholipids (membrane proteins). The remaining 24 (zinc-binding proteins, RNA-binding proteins, and nuclear body proteins) also undergo extensive phosphorylation. On average, these 24 proteins have one phosphosite for every ten residues, five times more than expected by random chance (**Fig. S10**). In short, we find for essentially every single example of an abundant, positively charged protein, that its charge is neutralized by a phosphate moiety—either in *trans* (nucleic acids or phospholipid headgroups) or in *cis* (through phosphorylation).

To examine the generality of this finding, we extended our analysis across five other model organisms (*A. thaliana*, *X. laevis*, *S. pombe*, *S. cerevisiae*, and *E. coli*). In all five cases, the same trends held true, with almost no examples of abundant free cytosolic positively charged proteins identified (**Fig. S11**, **Table S3**). Taken together, our integrative bioinformatic analysis suggests that while proteomes do possess many examples of proteins with a net positive charge, these are either expressed at low copy numbers, sequestered from the cellular environment via constitutive binding to anionic biomolecules, or neutralized through phosphorylation.

---

**Figure S1:**
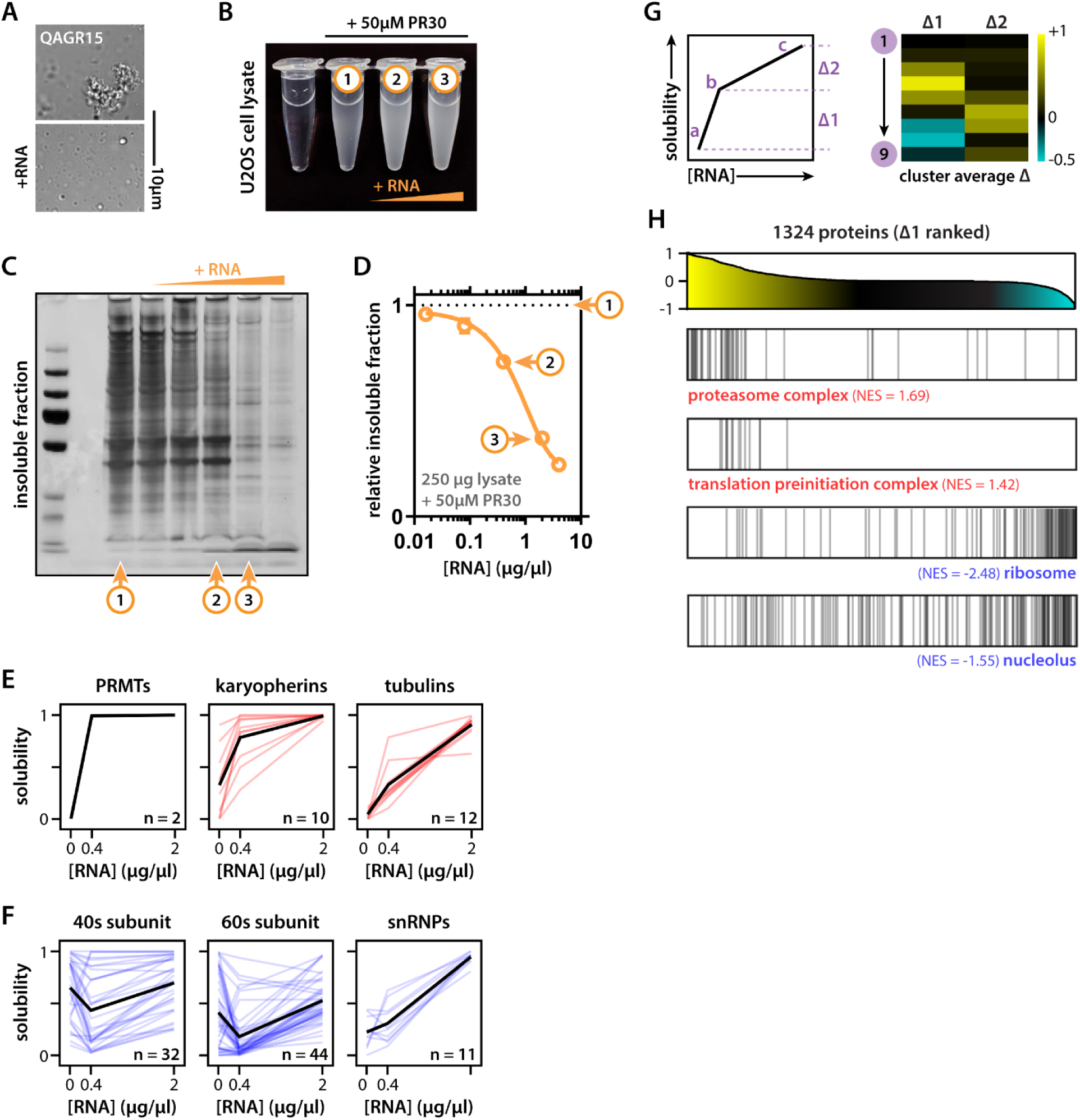
Disease-related basic repeat peptides target biomolecular condensates. (A) QAGR + lysate condensates are modulated by RNA addition (2 µg/µl). (B) PR + lysate mixtures with (1) no, (2) 0.4 and (3) 2µg/µl of RNA added. (C) Silver stain of condensate fraction highlighting position of the three mixtures. Same picture as in Fig. 1. (D) Quantification of total condensate fraction. (E-F) Example solubility profiles of red and blue protein families. Colored lines indicate individual proteins, black lines indicate family average. (G-H) Solubility profiles can be broken down in solubility steps. Δ1 indicates the first solubility step (i.e., response to the addition of 0.4 µg/µl RNA) and has opposite values for blue versus red clusters (convex vs concave profiles). Different protein classes enrich for high or low Δ1 values in our data set. GSEA analysis^137^, NES = normalized enrichment score.

**Figure S2:**
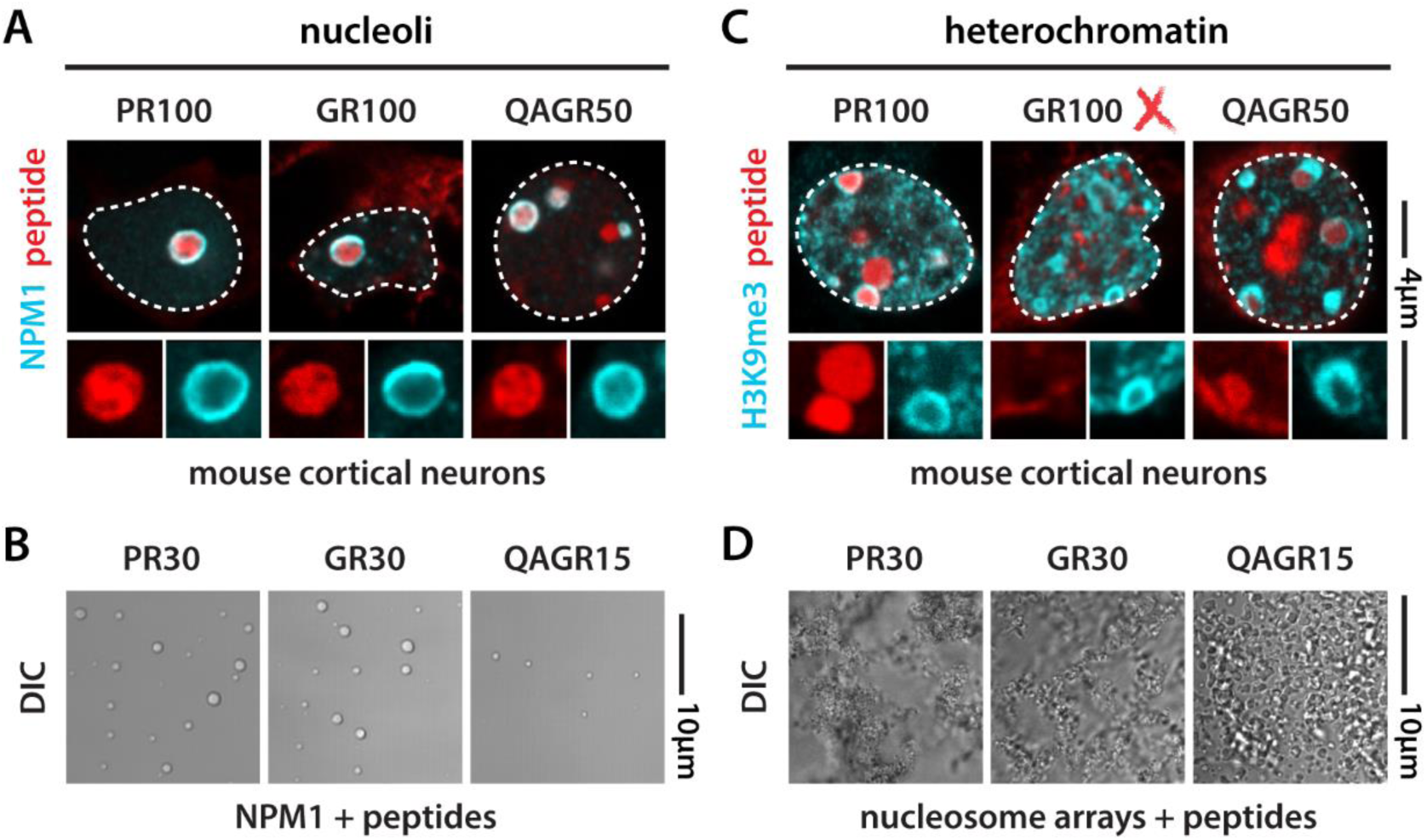
Disease-related basic repeat peptides target biomolecular condensates. (A) mCherry-tagged PR100, GR100 and QAGR50 target nucleoli in mouse primary cortical neurons. (B) PR30, GR30 and QAGR15 peptides phase separate with NPM1 *in vitro*. (C) mCherry-tagged PR100 and QAGR50, but not GR100, target heterochromatin in primary cortical neurons. (D) PR30, GR30 and QAGR15 peptides condense nucleosome arrays *in vitro*. This shows that despite GR can engage with chromatin in the test tube, in cells it seems to prefer other compartments.

**Figure S3:**
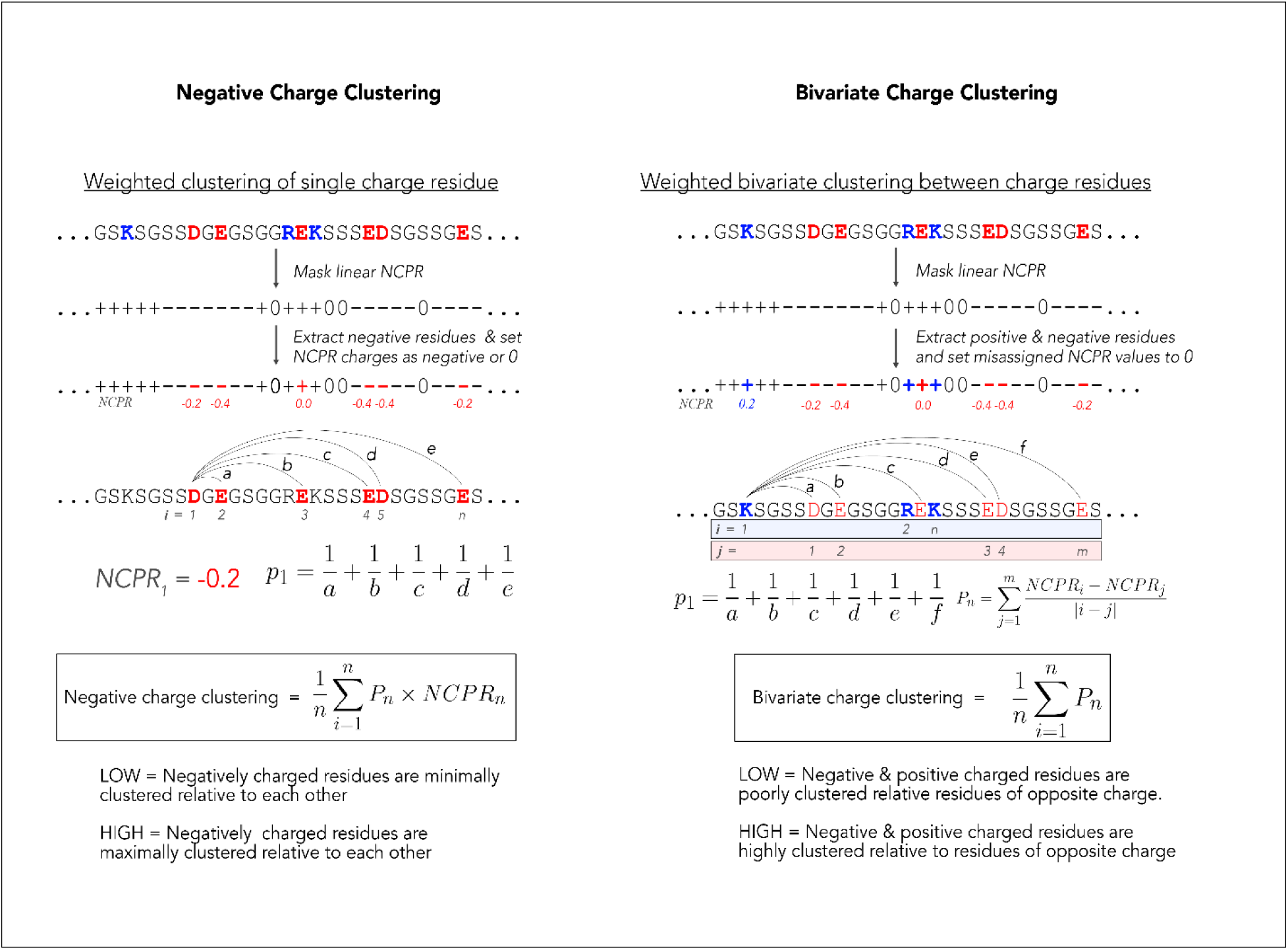
Overview of our computational approach for calculating charge clustering.

**Figure S4:**
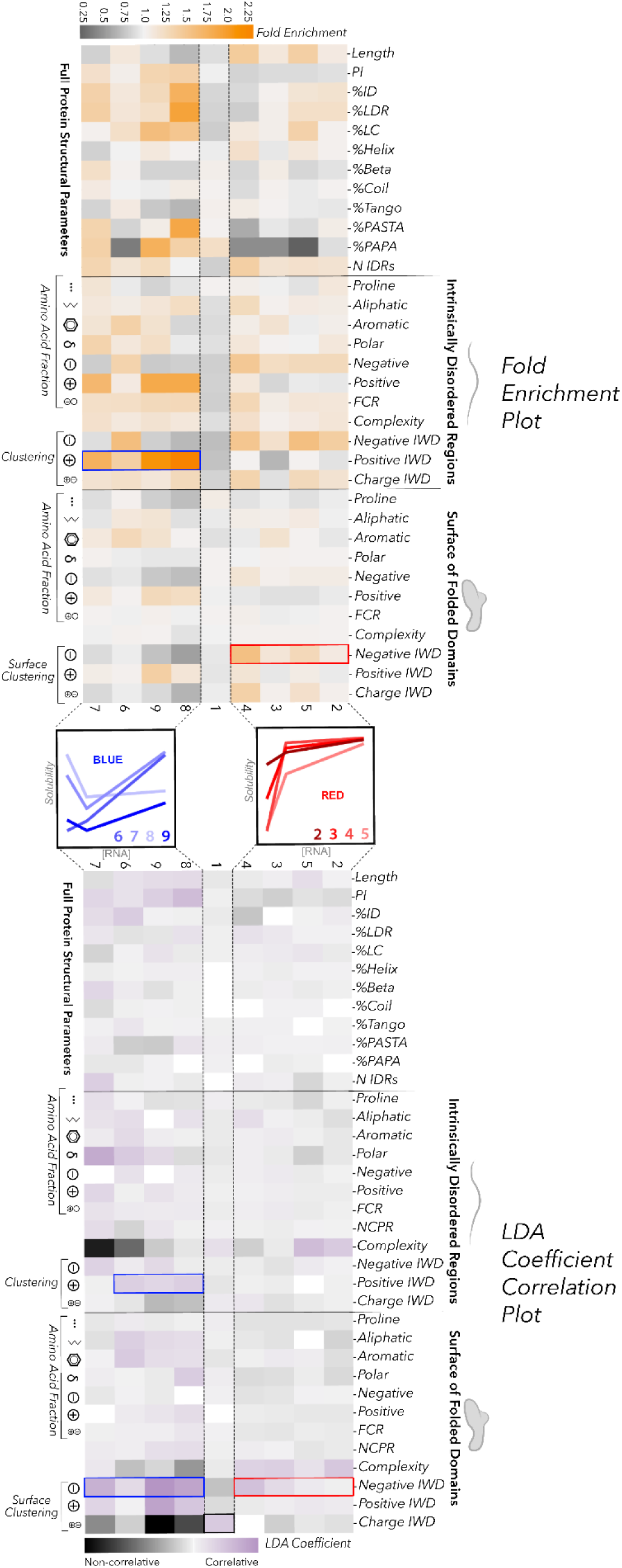
Enrichment and correlation of sequence features in different clusters. The upper heat map shows the fold enrichment of sequence features, while the bottom heatmap reports on the correlation of the sequence feature with the cluster (coefficients of linear discriminants as calculated from an LDA). In both heatmaps, each column denotes a cluster with blue clusters on the left and red clusters on the right, and each row denotes a different sequence feature. Sequence features are grouped by features calculated using residues in just the IDRs, features calculated based on residues on the surface of the folded domains, and features calculated on the entire protein sequence. The magnitude of the LDA coefficients in the bottom heatmap describe how well correlative a feature is with each cluster, while the sign of the coefficient denotes correlation (negative coefficients describe non-correlative features and do not inform on anti-correlation).

**Figure S5:**
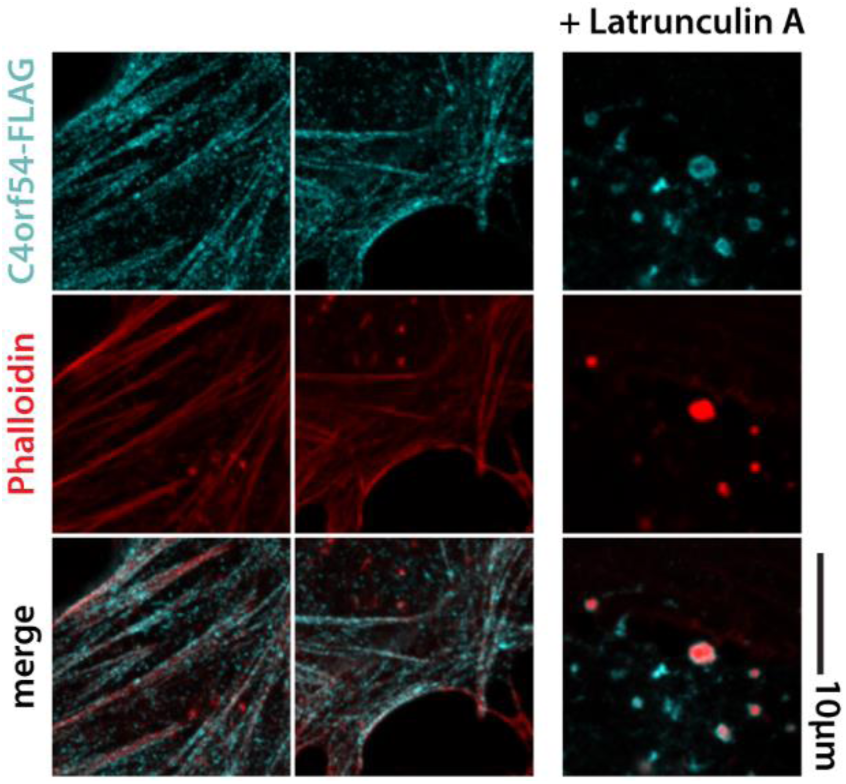
C4orf54 is an actin-binding protein. Expression of C4orf54-FLAG in U2OS cells shows colocalization with the actin cytoskeleton (phalloidin staining). C4orf54 forms shells around condensed actin after treatment with latrunculin A.

**Figure S6:**
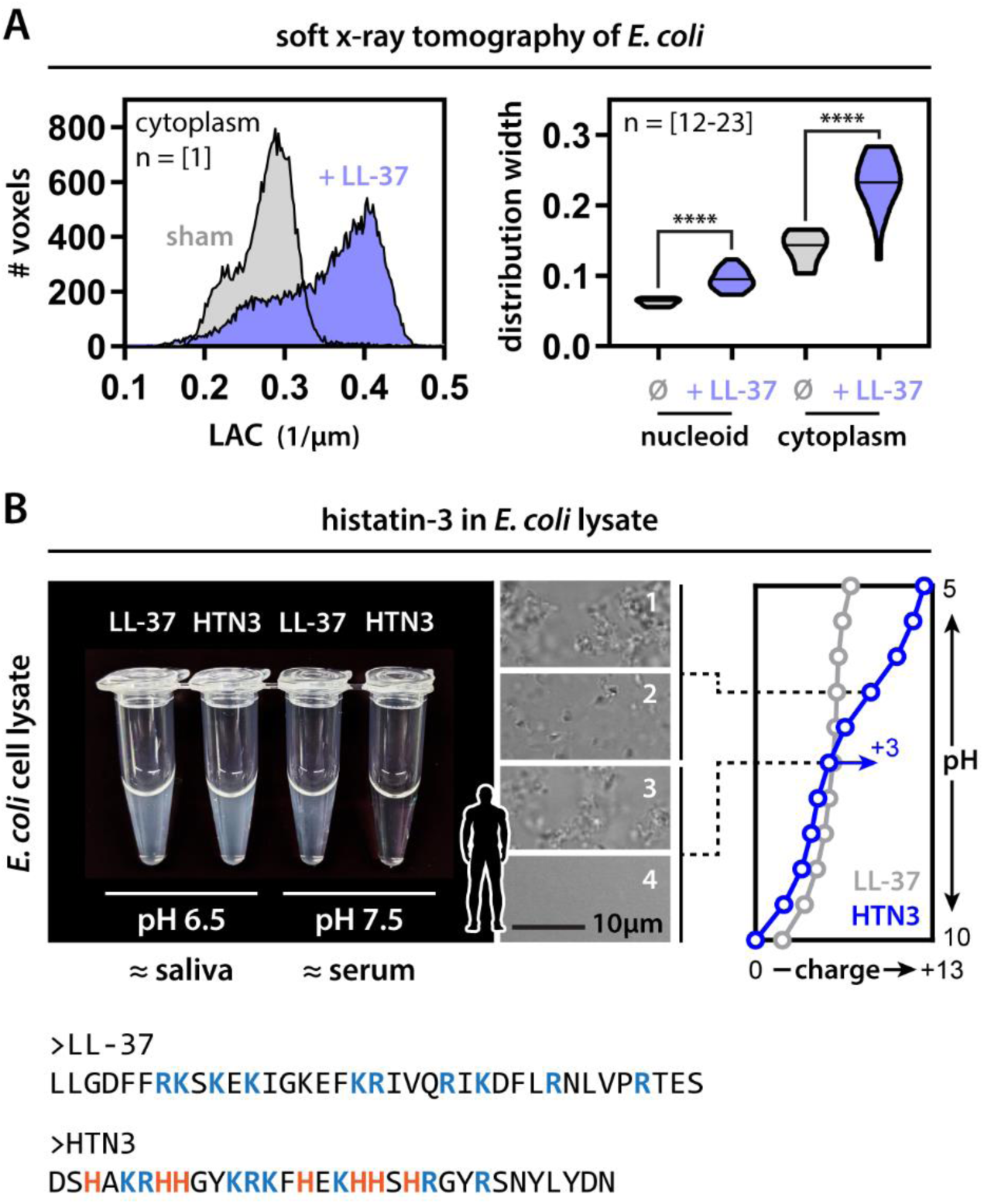
LL-37 and HTN3. (A) Molecular density (reported as the X-ray linear absorption coefficient or LAC) distribution of individual *E. coli* cells imaged with soft x-ray tomography shows increase in heterogeneity upon LL-37 treatment (i.e., wider distribution). This change in heterogeneity is observed for both nucleus and cytoplasm. Mann-Whitney. **** p-value < 0.0001. (B) HTN3 only drives *ex vivo* condensate formation under conditions similar to those of saliva (i.e., mildly acidic). Under these conditions HTN3 gains +3 net charge, while LL-37 retains the same charge.

**Figure S7:**
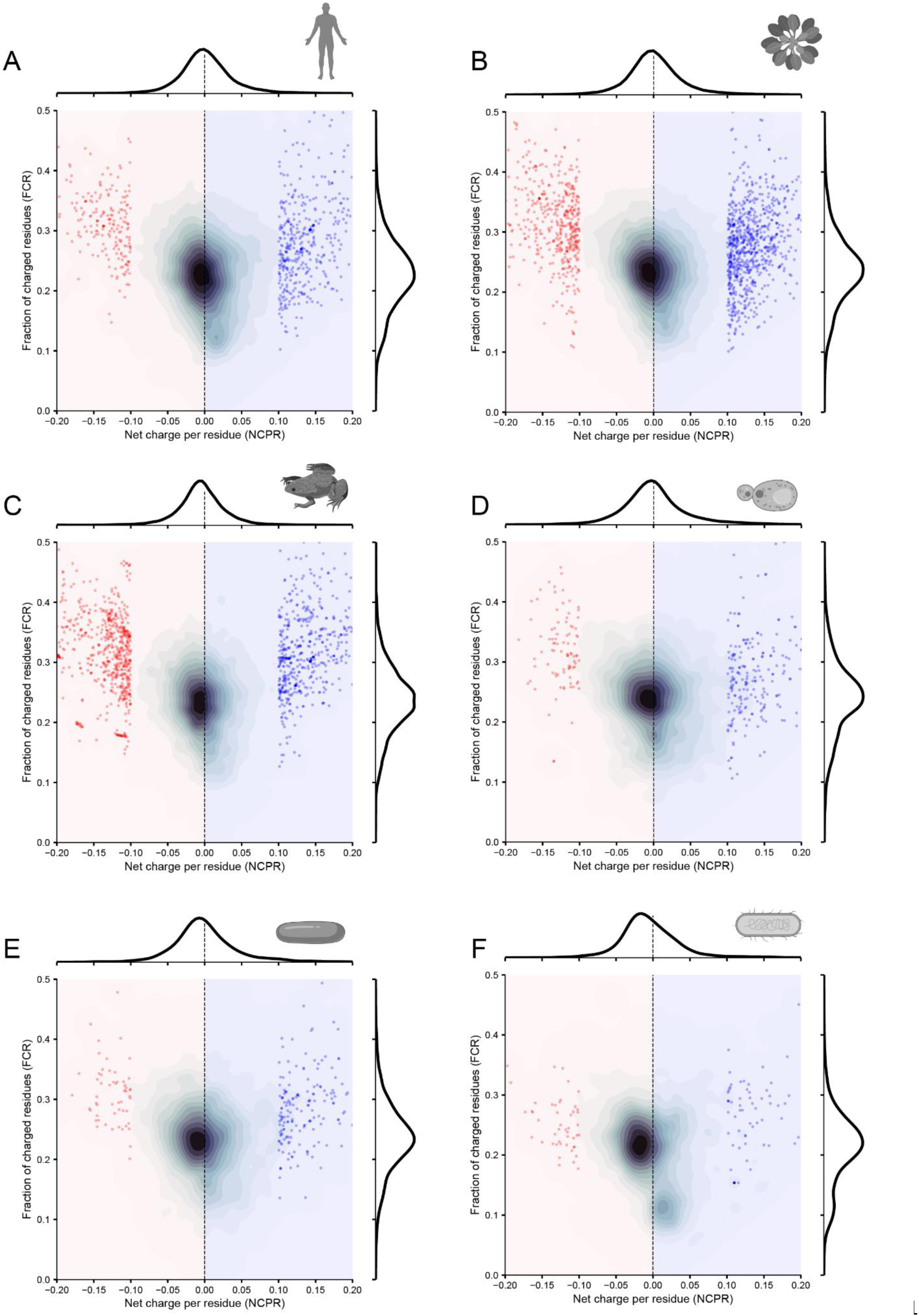
Proteome-wide analysis for six different organisms reveals a small bias towards negatively charged proteins, yet many examples of positively charged proteins exist. This analysis examined (A) *Homo sapiens* (n=20,393), (B) *Arabidopsis thaliana* (n=39,319), (C) *Xenopus laevis* (n=49,880), (D) *Saccharomyces cerevisiae* (n=6,060), (E) *Schizosccharomyces pombe* (n=5,122), and (F) *Escherichia coli* (n=4,438). In all cases, there are approximately an equal number of proteins with a net positive charge as a net negative charge. Proteins with a net charge per residue above 0.10 or below –0.10 are identified as individual markers, while all other proteins are reported as contour density data. Marginal distributions for the net charge per residue (NCPR) and fraction of charged residues (FCR) are shown alongside the two-dimensional distributions.

**Figure S8:**
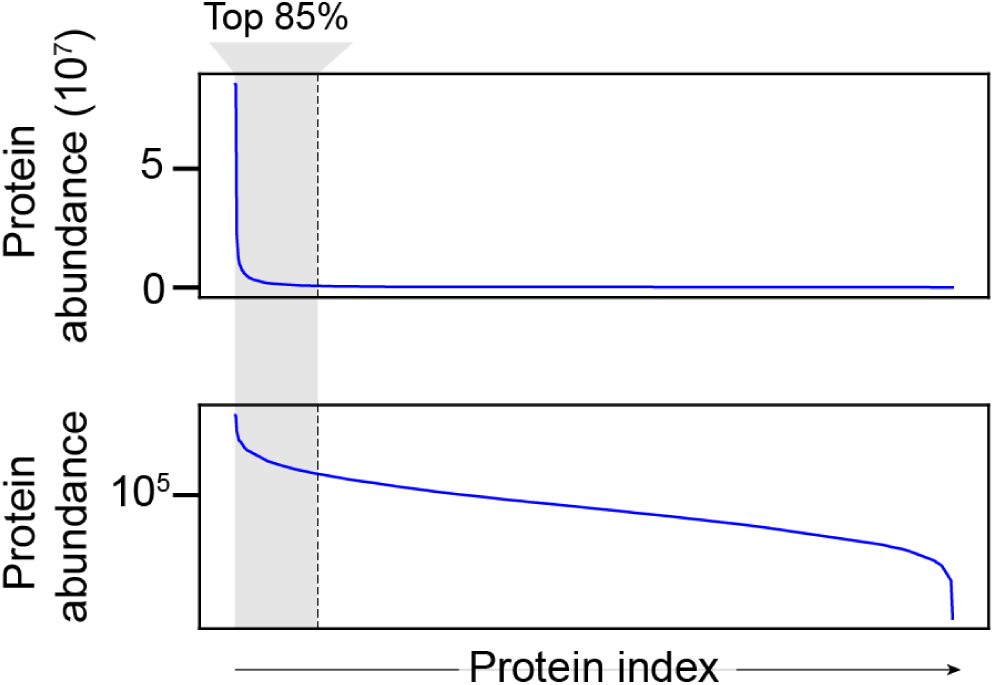
Quantitative mass-spectrometry data from Hein *et al.*^128^ provides copy number information for 9,209 proteins. We rank-ordered those proteins by most to least abundant and took the set of proteins that comprise the top 85% of proteins in a human cell by copy number. This ensures our analysis only focuses on a subset of proteins that are found in high abundance (i.e., in human cells, ∼10,000 copies or more). This same approach was used to select high-abundance proteins across five other organisms.

**Figure S9:**
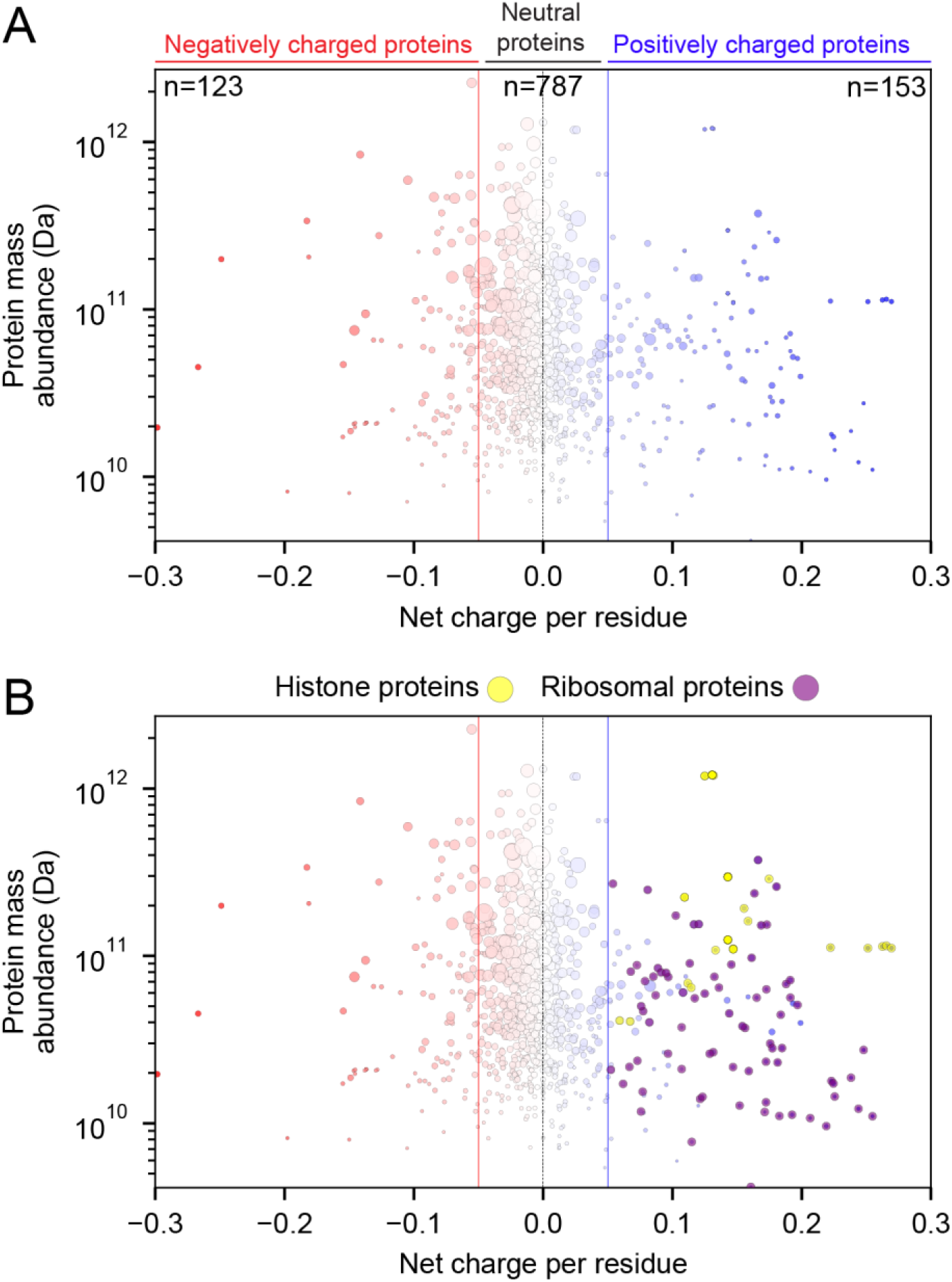
Assessment of highly abundant proteins across the human. (A) Of the 1063 highly abundant proteins identified, just 153 have a net positive charge, and of those, the majority (∼90%) are sequestered in constitutive molecular complexes with RNA (ribosomal proteins, RNP proteins), DNA (histones) or phospholipids (membrane proteins). The remaining proteins are nuclear RNA-binding proteins. (B) Same data as shown in panel A with histones and ribosomal proteins explicitly labelled for convenience.

**Figure S10:**
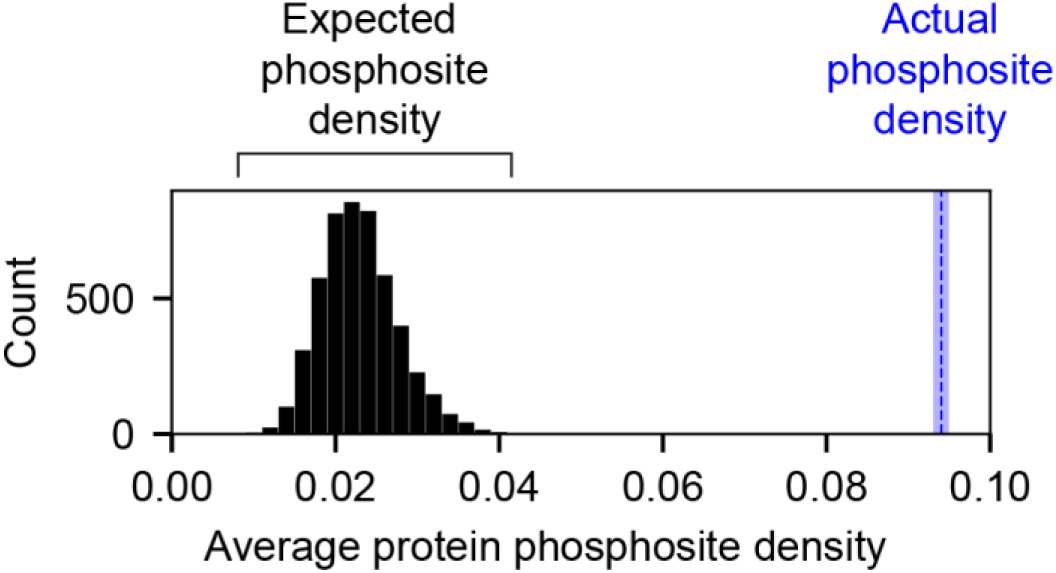
The expected phosphosite density was generated by randomly selecting 24 proteins and calculating the average phosphosite density (number of residues that possess phosphosites / total number of residues) 5000 times with replacement to construct a null distribution. This null distribution gives an expected average phosphosite density of 0.02 (i.e., one in every fifty residues is phosphorylated). The value and the associated distribution can be compared with the actual phosphosite density for the 24 non-RNP/membrane-bound proteins identified from our 153 highly abundant positively charged human proteins. The phosphosite density for these 24 proteins is 0.94, or one in every ten residues.

**Figure S11:**
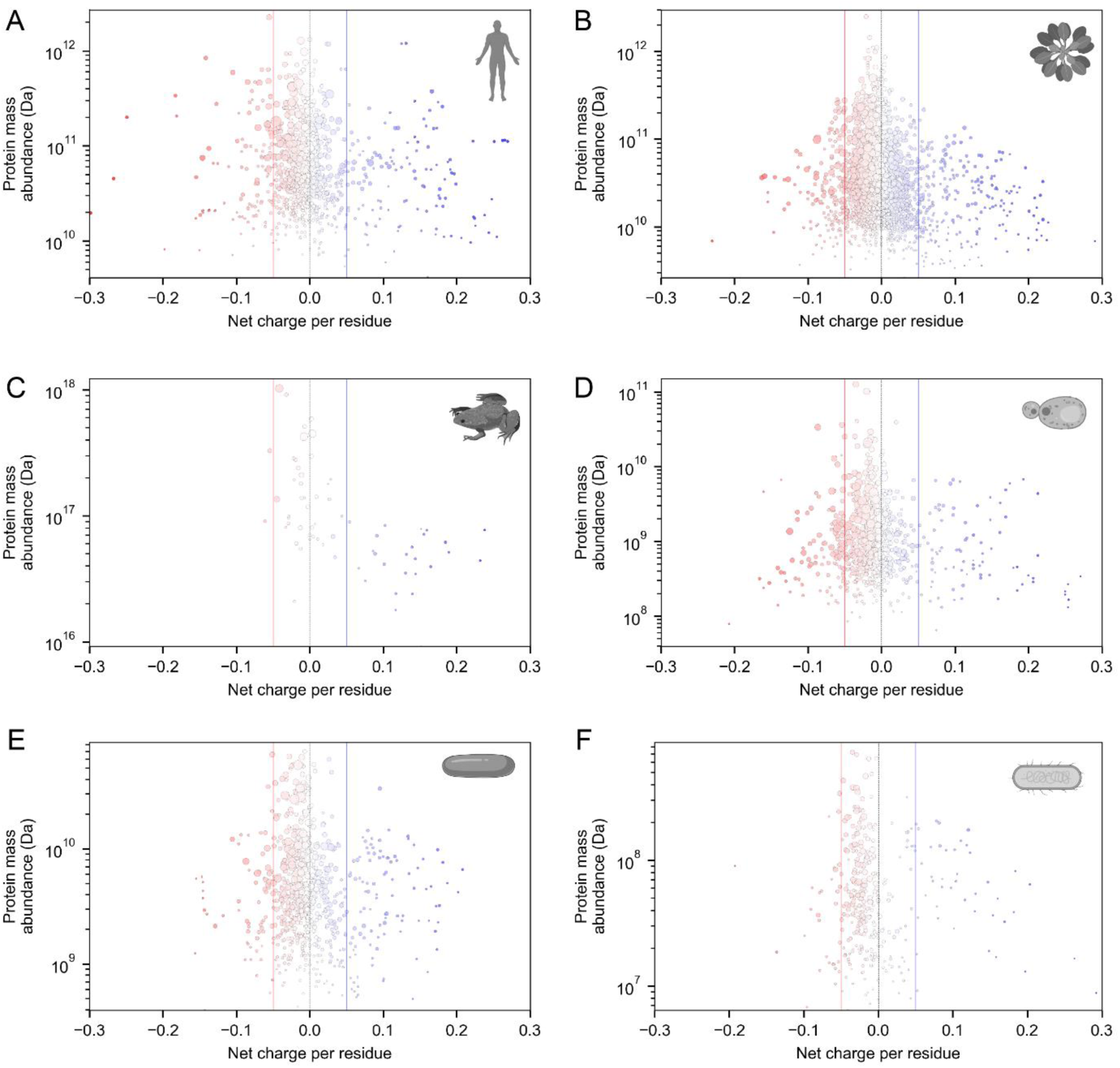
Analysis of abundant, positively charged proteins across the tree of life consistently identifies positively charged proteins as histones, ribosomal proteins, membrane proteins, or proteins engaged as parts of ribonuclear protein (RNP) complexes (see **Table S3**).

**Figure S12:**
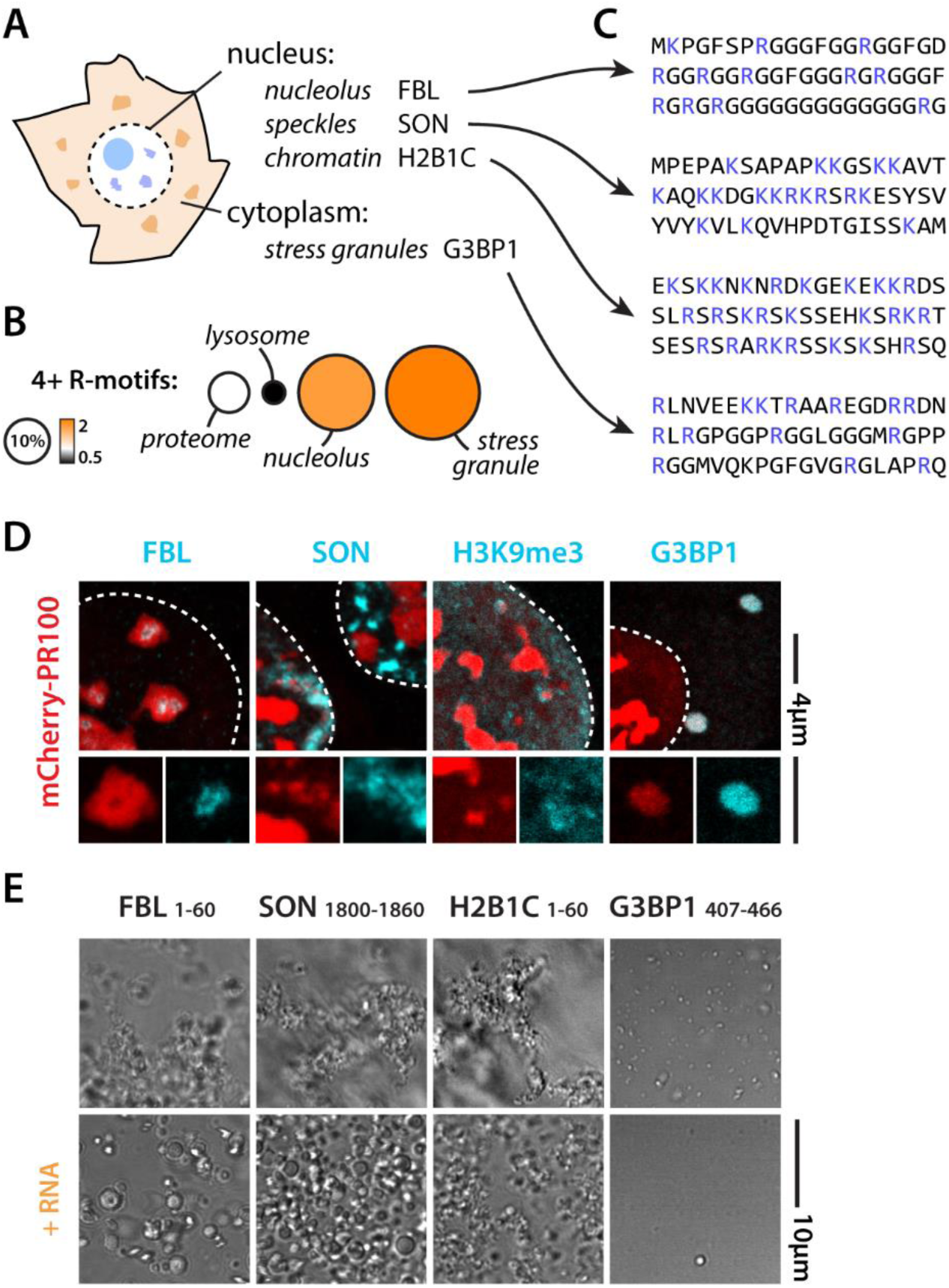
Endogenous cationic IDRs show similar behavior to disease ones when in isolation. (A) Key proteins from several RNA-centric condensates (B) RNA-centric condensates are enriched in arginine-rich motifs (fraction of proteins with at least four motifs^138^). Circle size indicated percentage of proteins with arginine-rich motif. Color indicated fold enrichment. (C) Example cationic IDRs from key condensate proteins. (D) These condensate proteins all colocalize with PR when expressed in cells. (E) Isolated cationic IDR peptides form irregular gel-like condensates when added to cell lysate, but are modulated into more spherical assemblies upon addition of RNA (2 µg/µl).

